# They all rock: A systematic comparison of conformational movements in LeuT-fold transporters

**DOI:** 10.1101/2024.01.24.577062

**Authors:** Jacob A. Licht, Samuel P. Berry, Michael A. Gutierrez, Rachelle Gaudet

## Abstract

Many membrane transporters share the LeuT fold—two five-helix repeats inverted across the membrane plane. Despite hundreds of structures, whether distinct conformational mechanisms are supported by the LeuT fold has not been systematically determined. After annotating published LeuT-fold structures, we analyzed distance difference matrices (DDMs) for nine proteins with multiple available conformations. We identified rigid bodies and relative movements of transmembrane helices (TMs) during distinct steps of the transport cycle. In all transporters the bundle (first two TMs of each repeat) rotates relative to the hash (third and fourth TMs). Motions of the arms (fifth TM) to close or open the intracellular and outer vestibules are common, as is a TM1a swing, with notable variations in the opening-closing motions of the outer vestibule. Our analyses suggest that LeuT-fold transporters layer distinct motions on a common bundle-hash rock and demonstrate that systematic analyses can provide new insights into large structural datasets.

## Introduction

The amino acid polyamine-organocation (APC) superfamily is the second largest superfamily of secondary active transporters (Pfam clan CL0062), is composed of 19 protein families (Table 1) and is ubiquitous in all kingdoms of life^1^. APC-superfamily transporters are specific for a variety of substrates, including sugars, amino acids, neurotransmitters, and metal cations. The transport of these substrates is often energetically unfavorable and is typically coupled to the cotransport of a cation, such as H^+^ or Na^+^, along a concentration gradient, a process known as secondary active transport. APC-superfamily transporters are present in all organisms and play important roles in human health, including in the immune response, digestion, neural signaling, and maintaining blood glucose levels. They are involved in diseases that affect a large percentage of the global population, including anemia, bipolar disorder, depression, diabetes, and immunodeficiency^1,2^.

**Table 1.** The 19 protein families of the APC superfamily.

| Abbreviation | Name | TC number | SLC family | Co-substrate(s) | Number of structures (homologs)<sup>a</sup> |
| --- | --- | --- | --- | --- | --- |
| APC | Amino Acid-Polyamine-Organocation | 2.A.3 | 7 |  | 45 (10) |
| BCCT | Betaine/Carnitine/Choline Transporter | 2.A.15 | - | Na<sup>+</sup> symport | 24 (3) |
| AAAP | Amino Acid/Auxin Permease | 2.A.18 | 32, 36, 38 |  | 4 (1) |
| SSS | Solute:Sodium Symporter | 2.A.21 | 5 | Na<sup>+</sup> symport | 8 (2) |
| NSS | Neurotransmitter:Sodium Symporter | 2.A.22 | 6 | Na<sup>+</sup> symport | 141 (6) |
| AGCS | Alanine or Glycine:Cation Symporter | 2.A.25 |  | Na<sup>+</sup> or H<sup>+</sup> symport | 4 (1) |
| LIVCS<sup>b</sup> | Branched Chain Amino Acid:Cation Symporter | 2.A.26 |  | Na<sup>+</sup> or H<sup>+</sup> symport |  |
| CCC | Cation-Chloride Cotransporter | 2.A.30 | 12 |  | 45 (8) |
| NCS1 | Nucleobase:Cation Symporter-1 | 2.A.39 |  |  | 6 (1) |
| HAAAP<sup>b</sup> | Hydroxy/Aromatic Amino Acid Permease | 2.A.42 |  |  |  |
| Nramp | Metal Ion (Mn<sup>2+</sup> - Fe<sup>2+</sup>) Transporter | 2.A.55 | 11 | H<sup>+</sup> symport | 19 (3) |
| KUP | K<sup>+</sup> Uptake Permease | 2.A.72 |  | H<sup>+</sup> symport | 2 (1) |
| CstA<sup>b</sup> | Putative Peptide Transporter Carbon Starvation | 2.A.114 |  |  |  |
|  | CstA |  |  |  |  |
| PAAP<sup>b</sup> | Putative Amino Acid Permease | 2.A.120 |  |  |  |
| <sup>b</sup> | TMEM104<sup>c</sup> |  |  |  |  |
| AE<sup>d</sup> | Anion Exchanger | 2.A.31 | 4 | Cl<sup>-</sup> antiport or Na<sup>+</sup> symport |  |
| NCS2<sup>d</sup> | Nucleobase:Cation Symporter-2 | 2.A.40 | 23 |  |  |
| BenE<sup>d</sup> | Benzoate:H<sup>+</sup> Symporter | 2.A.46 |  | H<sup>+</sup> symport |  |
| Sul<sup>d</sup> | Sulfate Permease | 2.A.53 | 26 |  |  |

<sup>a</sup>Each independent LeuT-fold chain in a PDB entry is counted as a structure, and the number of homologs includes both paralogs and orthologs. Only structures of LeuT-fold families available in the PDB as of June 11, 2021, are catalogued.
<sup>b</sup>Although no structures are available, these families are predicted to adopt a LeuT fold, including in predicted structures in the AlphaFold Database (Example members: 2.A.26, *E. coli* BrnQ, Uniprot ID P0AD99; 2.A.42, *E. coli* TyrP, Uniprot ID P0AAD4; 2.A.114, *E. coli* CstA, Uniprot ID P15078; 2.A.114, *E. coli* CstA, Uniprot ID P15078; 2.A.120, *B. subtilis* YyaD, Uniprot ID P37520; Human TMEM104, Uniprot ID Q8NE00)<sup>30</sup>.
<sup>c</sup>TMEM104 is a human protein recently classified as an APC but without TC or SLC classification yet<sup>31</sup>.
<sup>d</sup>These four families are classified within the APC superfamily but have either experimentally determined or predicted UraA folds<sup>1,30,32</sup>.

Our understanding of the transport mechanisms of APC-superfamily transporters has been greatly enhanced by the determination of hundreds of high-resolution structures over the past twenty years. These structures revealed that despite poor pairwise sequence identity, most APC-superfamily transporters share close structural homology^3^. Fifteen of the 19 families share a common fold, known as the LeuT fold, which was first identified from a crystal structure of *Aquifex aeolicus* LeuT, a bacterial Na^+^-coupled amino acid importer (Table 1)^4^. The LeuT fold comprises two repeats of 5 transmembrane (TM) helices inverted with respect to the membrane plane^5^. Although many transporters have additional TMs N- or C-terminal to the LeuT fold, here we refer to TMs of all discussed transporters by their placement within the LeuT-fold core. The first two TMs of each repeat (TMs 1, 2, 6, 7) form the bundle, and the last three TMs of each repeat form the scaffold, which is subdivided into the hash (TMs 3, 4, 8, 9) and arms (TM5 and TM10) (Figure 1A). Portions of the first and third TM of each inverted repeat (TMs 1, 3, 6, 8) compose a central binding site shaped to accommodate the transporter’s specific primary substrate and ion co-substrate pair^6^.

**Figure 1.**
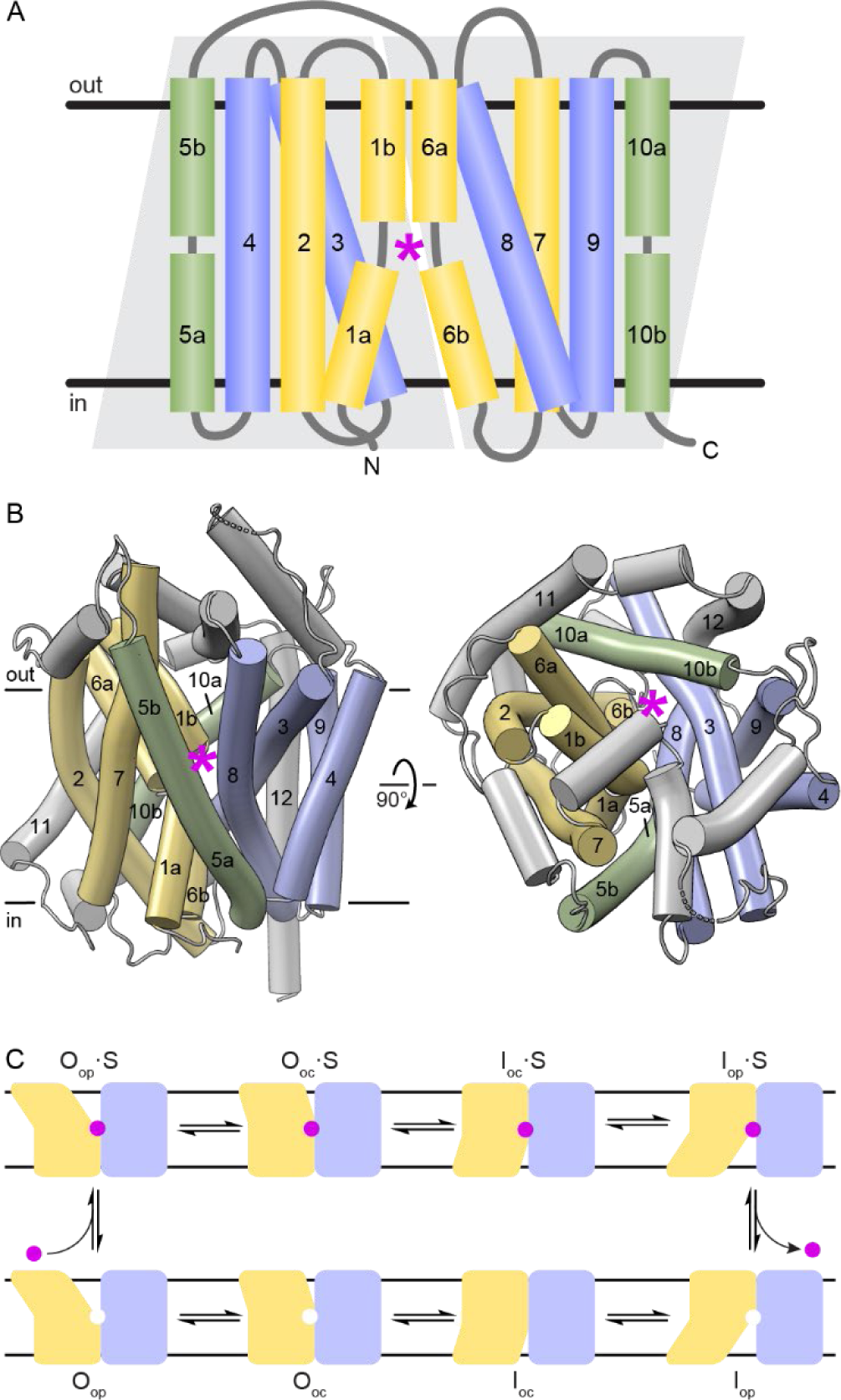
The LeuT-fold superfamily of secondary active transporters. (A) Topology of the 10 LeuT-fold core TM helices in the membrane. Light grey trapezoids delineate the two five-helix inverted repeats. TM helices are colored as follows: bundle in yellow, hash in blue, arms in green. The general position of the substrate-binding site is indicated by a magenta star. Any non-core structural elements like the termini and links between TM helices are in grey. These vary considerably in length between homologs, in some cases including additional N- or C-terminal TM helices and/or extracellular or intracellular helices. The same color scheme is used in later figures. (B) Structure of LeuT (PDB: 2A65), viewed from the membrane plane (left) or the extracellular side (right) and colored as in panel A. LeuT has two additional C-terminal TM helices beyond the LeuT-fold core, labeled 11 and 12. (C) LeuT-fold transporters cycle between different conformational states. The conformations illustrated from left to right are: O_op_ – outward open; O_oc_ – outward occluded; I_oc_ – inward occluded; I_op_ – inward open. Stable intermediate O_oc_ and I_oc_ conformations may not exist for all LeuT-fold transporters. The top row contains substrate (S) bound conformations, and the bottom row the corresponding substrate-free (or antiported substrate bound, depending on the transporter) conformations.

Substrates access the binding site through alternating access from the outer and intracellular aqueous vestibules; outer vestibule closing is coupled to intracellular vestibule opening and vice versa, so both vestibules cannot simultaneously be open^7,8^. As most LeuT-fold transporters are importers, they bind their primary substrate in an outward-open (O_op_) conformation, transition through one or two occluded intermediates—outward-occluded (O_oc_) or inward-occluded (I_oc_)— and release it in an inward-open (I_op_) conformation (Figure 1B). We define a conformation as outward (O) or inward (I) depending on whether the outer vestibule or intracellular (cytosolic) vestibule is solvated, respectively; we define a conformation as occluded if the binding site is occluded from bulk solvent and as open if the binding site is solvated. In other words, outward (O) and inward (I) are general terms to refer to either O_op_ or O_oc_, and either I_oc_ or I_op_, respectively.

Determining the degree to which variation in conformational movements is supported by the LeuT fold is important to understand how LeuT-fold transporters achieve such varied function across paralogs with vastly divergent substrates. A null hypothesis is that all LeuT-fold transporters share a universal mechanism of transport. However, the notion of a universal conformational cycle may not hold true across the entire LeuT-fold superfamily. The first structure of LeuT led to the suggestion that alternating access is achieved through gating networks of the intracellular and outer vestibules, with TM helices acting as gates and the unwound regions in TM1 and TM6 as the “joints” that allow for movement of the gates^4^. This model was followed by the rocking-bundle model of transport, which suggested that alternating access is achieved by rigid-body movement of the bundle relative to the scaffold^7^. While a rocking-bundle motion accounts for much of the outward-to-inward transition of LeuT-fold transporters, for multiple transporters the mechanisms are more complex or a combination of the two models^3,9,10^.

Anecdotal differences in the conformational cycle of LeuT-fold transporters have been described. For example, it has been reported that in the bacterial betaine transporter BetP the position of the bundle differs less between O_op_ and I_op_ than in other transporters^11^. The authors hypothesize that this conformational adaptation of BetP assists osmotic sensing by keeping the substrate pathway less solvated, suggesting that these differences in conformational movements could be functionally important.

However, it is unclear whether descriptions of the conformational transitions of LeuT-fold transporters differ because of intrinsic differences in the proteins or because we lack a systematic way to make comparisons. Applying a common computational pipeline to analyze a broad range of LeuT-fold transporters by leveraging the existing wealth of published structures of LeuT-fold transporters would allow a more systematic comparison of conformational differences. To elucidate how much variation is supported, we have undertaken a systematic comparison of the conformational movements for nine LeuT-fold transporters for which structures have been determined in multiple conformations. We use alignment-independent distance difference matrices (DDMs) to identify movements that are common to multiple transporters or unique to specific transporters. We find that all seven LeuT-fold transporters with experimentally determined structures in both outward and inward states show some extent of bundle-hash rocking, which, along with additional motions of TM1a and TM5a and the symmetry-related TM6a and TM10a, explain most of the outward-to-inward conformational changes.

## RESULTS

### Assembling a dataset of available LeuT-fold transporter structures

We created a comprehensive list of LeuT-fold structures published in the Protein Data Bank (PDB) as of June 11, 2021 using DALI^12^, yielding 298 unique chains (herein referred to as “structures”) from 28 unique proteins, some with more than one ortholog (Table 2). These proteins represent ten of the 15 LeuT-fold families (Table 1) and include six antiporters, two uniporters, and 20 symporters. Five transporters are H^+^-coupled, twelve are Na^+^-coupled, four are K^+^-coupled, one is coupled to both Na^+^ and K^+^, and three are uncoupled.

**Table 2.**
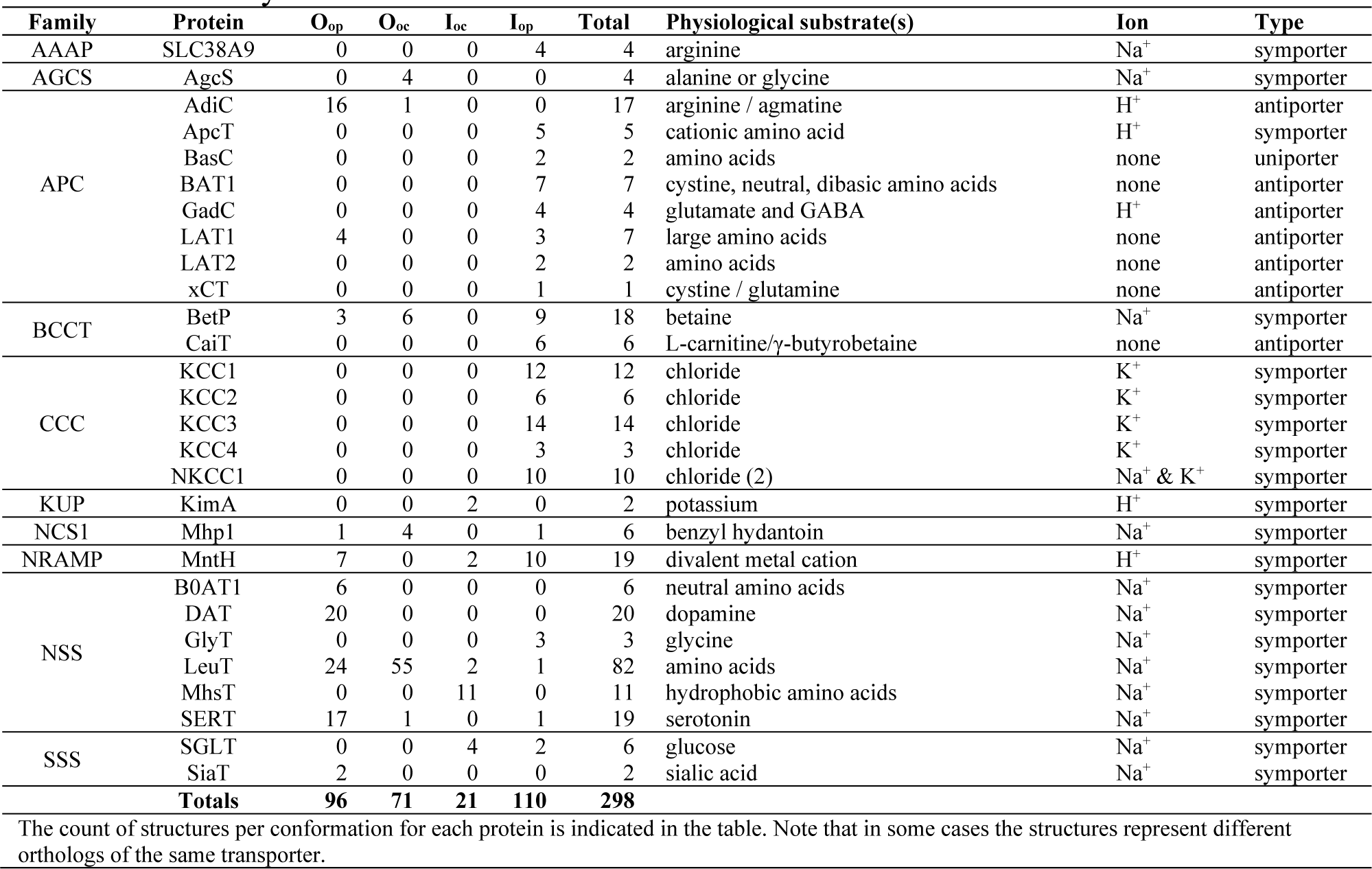
Summary of structures identified in the DALI search.

To efficiently annotate the conformation of each structure, we calculated pairwise root-mean-squared distance (RMSD) values for proteins with multiple structures and applied k-means clustering to the corresponding RMSD matrices (Figures S1 and S2). We annotated the conformation corresponding to each cluster by reviewing the corresponding literature (Supplementary Note and Tables S1 and S2). Of the 298 structures, 96 are O_op_, 71 O_oc_, 21 I_oc_, and 110 I_op_.

We note that for our analyses below, we assume that the conformational movements associated with substrate translocation are opposite and similar to those associated with the return to the starting state of the conformational cycle at the level of resolution required for the comparisons between different LeuT-fold transporters. This assumption is supported by the generally low RMSD values between multiple structures of a given conformation for each of the proteins in our dataset, regardless of whether or not they are substrate-bound (Figure S1, Table 3).

**Table 3.**
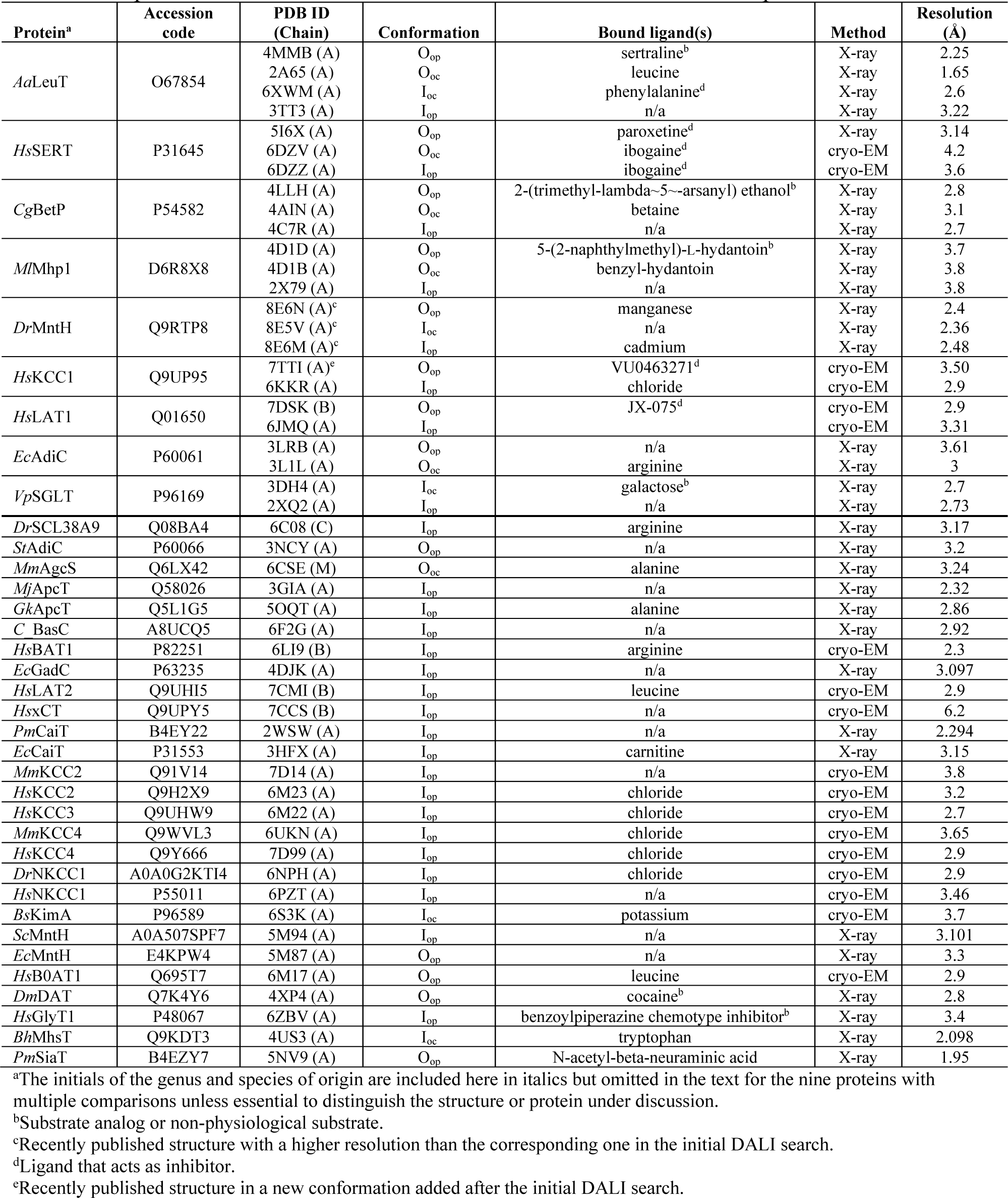
Representative structure selected for each conformation for each protein.

### Nine proteins have multiple conformations available

For each protein for which at least one structure has been determined, we chose one representative structure (generally the one with highest resolution) for each available conformation (Table 3). We also added a structure of human KCC1 determined in an additional conformation^13^ after we built our list and updated structures of *Deinococcus radiodurans* MntH to ones recently determined at higher resolution^14^. Nine proteins have structures available in multiple conformations (Table 3 and Table S3). Of the seven proteins with both O_op_ and I_op_ states available, only LeuT has two occluded states (O_oc_ and I_oc_). BetP, SERT, and Mhp1 have O_oc_, while MntH has I_oc_, and KCC1 and LAT1 have no available occluded state. Because several of these proteins have been extensively investigated, these results suggest that some transporters have only one or no occluded state as a well-populated intermediate^10,14^. In addition to the seven proteins listed above, AdiC structures are available for both outward conformations (O_op_ and O_oc_), and SGLT structures for both inward conformations (I_oc_ and I_op_). The structures of these nine LeuT-fold transporters provide a total of 22 pairwise comparisons between two conformations of the same protein, and the potential to compare the differences between two conformational states for at least two proteins except for O_oc_→I_oc_, for which only LeuT has structures.

### DDMs help define the conformational movements of the LeuT-fold core

We developed a methodology to quantify and compare the conformational changes between distant structural homologs like LeuT-fold transporters. Because the sequence similarity is very low across the LeuT-fold superfamily—including variations in the lengths of helices and loops— the sequences and structures of LeuT-fold transporters cannot be readily aligned with high confidence at residue-level resolution. Our strategy was therefore to develop a comparison using coarse-grained information and to focus our analysis on the internal motions of the ten conserved LeuT-fold TM helices. As detailed below, we devised a representation of the conformational differences observed between different conformations of a given transporter (Figure 2). With this representation, the same transition can readily be compared across different LeuT-fold proteins.

**Figure 2.**
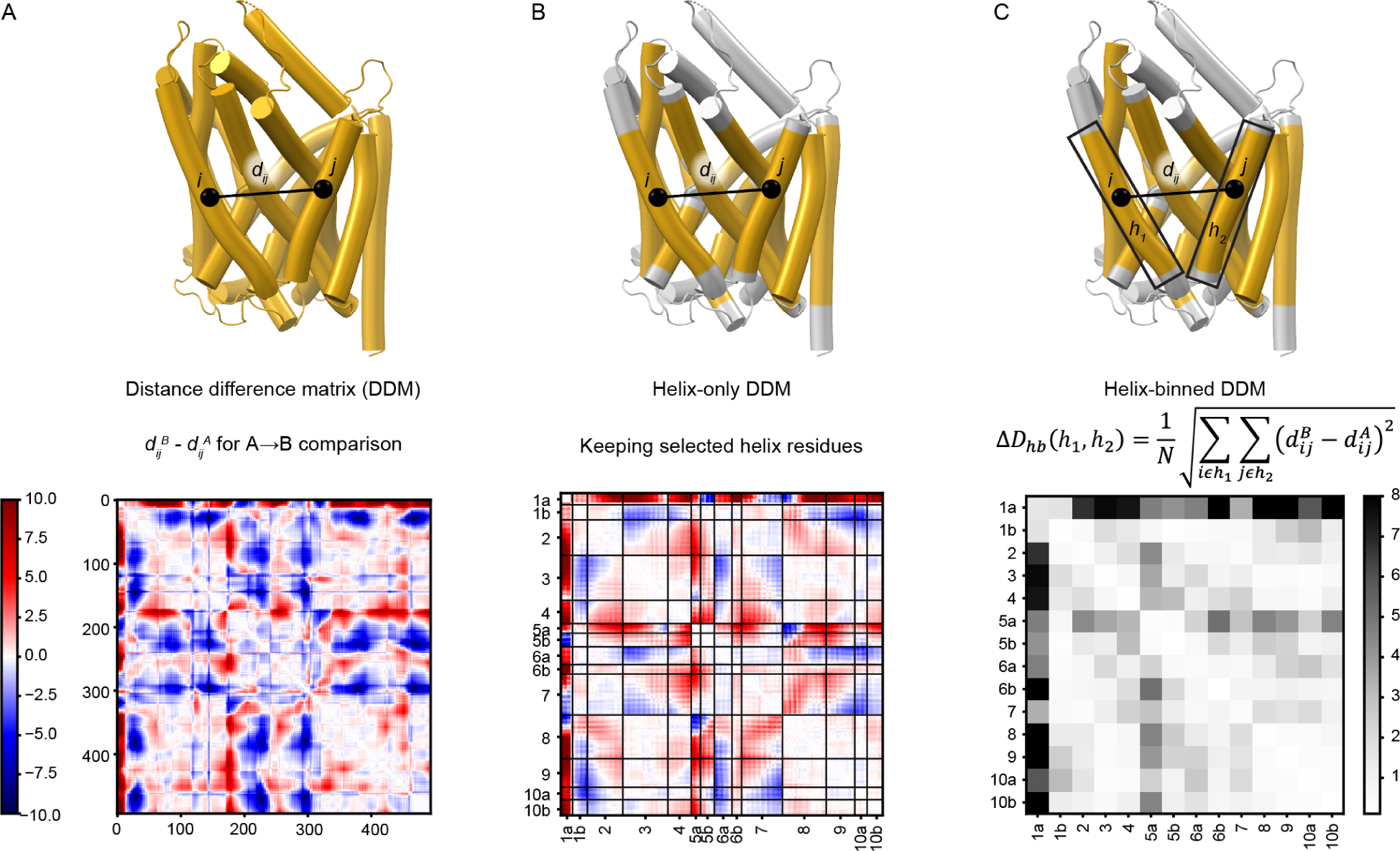
Distance difference matrices provide a visualization of conformational transitions. (A) The distance difference matrix comparing the O_op_ and I_op_ conformations of LeuT, with positions closer in the I_op_ conformation shown in red and those closer in the O_op_ conformation shown in blue. (B) A helix-only DDM (ho-DDM) for the same transition highlights only positions within the core transmembrane helices, the boundaries for which are highlighted in yellow on the structure above. The 14 helical segments of the LeuT fold are indicated on the axes. (C) A helix-binned DDM (hb-DDM) summarizes the information of each submatrix of the ho-DDM as a single value by taking the root-mean square of all distance difference values between each helix pair., generating an RMSDD value. The equation is fully described in the methods.

We first calculated a distance difference matrix (DDM; Figure 2A) for each of the 22 pairwise protein conformation comparisons for the nine LeuT-fold transporters with structures in multiple conformations by subtracting one structure’s distance matrix (distances between all pairs of Cα atoms) from that of the other^15,16^. An advantage of DDMs is that they rely solely on the internal coordinates of each of the two compared structures and thus avoid a potential perspective bias that can be introduced by a structural superposition, which requires choosing which structural elements to use for the superposition. The 22 DDMs represent six comparisons (Figure S3 and Table S3): O_op_→O_oc_ (5 proteins), O_op_→I_oc_ (2), O_op_→I_op_ (7), O_oc_→I_oc_ (1), O_oc_→I_op_ (4), and I_oc_→I_op_ (3). A representative DDM for the LeuT O_op_→I_op_ transition (Figure 2A) highlights regions that move toward (negative; blue) or away (positive; red) from one another, with static regions in white. As expected, many of these static regions fall near the diagonal, but others are off diagonal, indicating the presence of structural elements distant in the primary sequence that form rigid bodies in this transition.

To focus on the TM helices, we next built helix-only DDMs (ho-DDMs; Figures 2B and S3), which only include the DDM values for the residues within the core 10 LeuT-fold TMs. Because the middle of TM1 and TM6 include substrate-binding unwound regions and TM5 and TM10 are frequently kinked, we broke each into two segments (a and b), generating a set of 14 helical segments. The helix boundaries we used are listed in Table S4 and illustrated in Figure S4. For the LeuT O_op_→I_op_ transition (Figure 2B), the ho-DDM highlights how TM1a moves away from nearly all other helices as it swings outward from the protein, while other helices shift relative to some but not others. For example, the hash helices 3, 4, 8 and 9 move little relative to one another but have substantial motions relative to TMs 1, 2, 5, 6, and 7.

Finally, we built helix-binned DDMs (hb-DDMs; Figures 2C and S3)^17^, in which the submatrix of residue-by-residue values between each TM pair is converted into a root-mean-squared distance difference (RMSDD), yielding a 14-by-14 hb-DDM for the LeuT fold. By collapsing data into one value for each helix pair, hb-DDMs are uniformly sized for all LeuT-fold transporters, allowing for easy comparison both visually and quantitatively. The LeuT O_op_→I_op_ hb-DDM provides a simpler fingerprint of this particular transition, highlighting the large motions of TM1a and TM5a as well as a subtler pattern of interactions between other helices (Figure 2C).

To validate our choice of 14 helical segments, we extracted the values along the diagonal from all 22 hb-DDMs, representing intrahelix RMSDD values: a low value indicates that the helix itself is rigid, with minimal flexing between the two protein conformations under comparison. The average intrahelix RMSDD was 0.40 Å (standard deviation of 0.32 Å), in comparison to an average interhelix RMSDD of 1.5 Å (standard deviation of 1.5 Å). Furthermore, every helix has a distribution of intrahelix RMSDD values much more similar to the other intrahelical RMSDD values than to the distribution of interhelical RMSDD values (Figure S5 and Table S6).

### LeuT-fold transporters have a conserved rigid hash but variably flexible bundles

We first used this dataset of hb-DDMs to identify which groups of helices behave as rigid bodies across the transport cycle. We performed hierarchical clustering of the helices for each of the seven available O_op_→I_op_ hb-DDMs (Figures 3A-3C and S6). The hash helices (TMs 3, 4, 8 and 9) form a tight cluster for all proteins (except for TM8 in BetP; Figure S6), sometimes along with arm helices such as TM5a (BetP), TM5b (LAT1), or TM10a and b (LeuT and SERT; Figure 3A). By contrast, the traditional bundle of TMs 1, 2, 6, and 7 forms a tight cluster in some transporters such as Mhp1 (Figure 3B), KCC1, and BetP, but less so in others such as MntH (Figure 3C) or LeuT (Figure 3A), in which certain bundle helices move independently from one another. Arm helices TM10a and TM10b often cluster with the hash, while TM 5a and 5b often cluster with the bundle (Figures 3A-3C and S6). Overall, this hierarchical clustering analysis suggests that the hash serves as a rigid body across the LeuT-fold superfamily, while the bundle does not. In our illustrations, we superimpose structures using the rigid hash to better highlight the differences between conformations.

**Figure 3.**
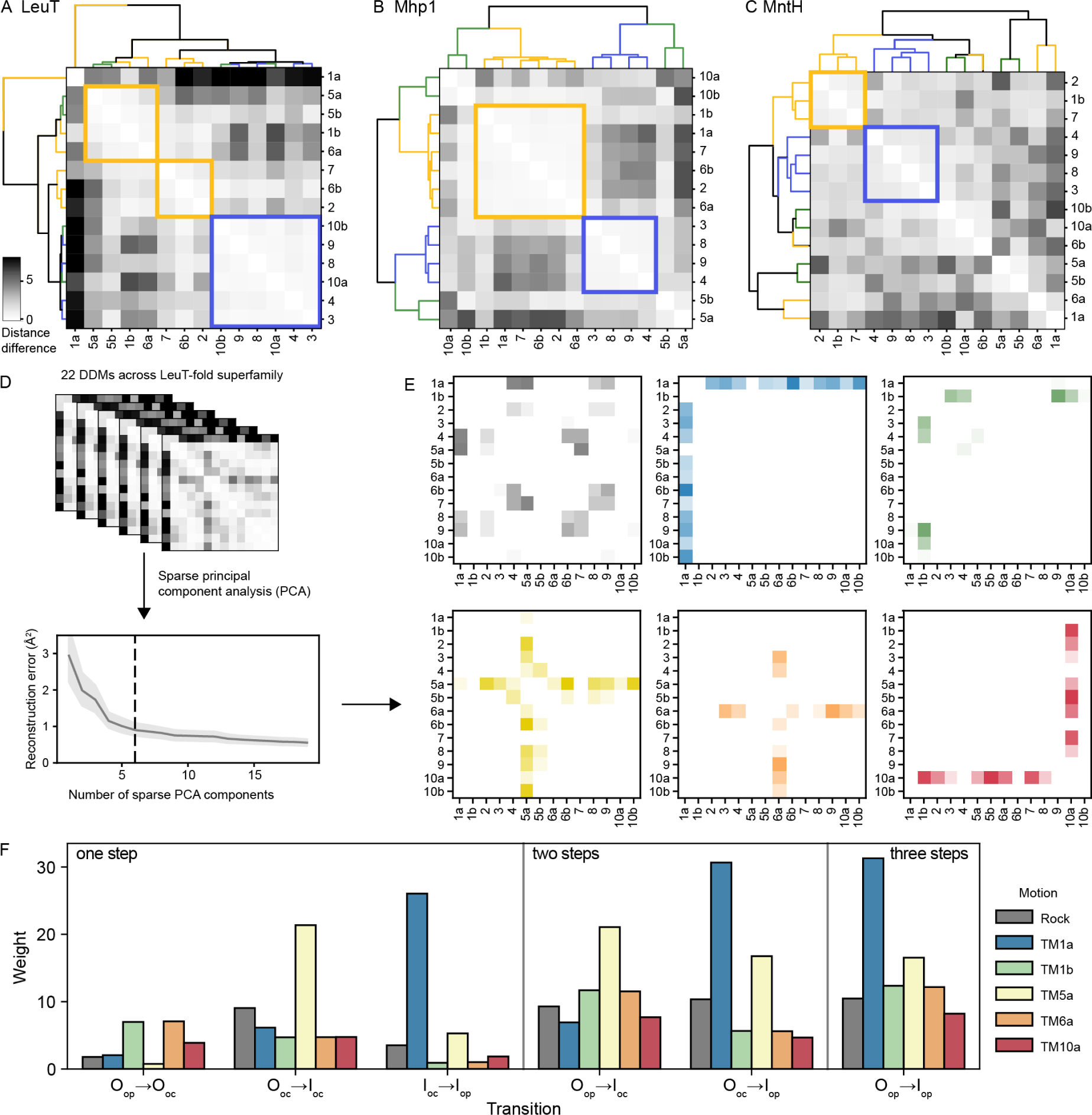
Clustering and dimensionality reduction on hb-DDMs provide new windows into LeuT-fold superfamily conformational change mechanisms. (A-C) Using the hb-DDMs to cluster the 14 helix segments highlights rigid bodies, with helix clusters most closely matching the bundle and hash represented in yellow and blue, respectively. hb-DDMs are shown for the O_op_→I_op_ transition of LeuT, Mhp1, and MntH. Dendrograms are colored according to canonical assignments of hash, bundle, and arms in yellow, blue, and green, matching Figure 1A. (D) Sparse PCA on the set of 22 hb-DDMs reveals at least six principal motions; while the mean square reconstruction error of sparse PCA decreases with increasing numbers of components, the improvement begins to level off after six sparse PCs. (E) The 6 sparse PCs, represented as 14-by-14 matrices, represent six distinct motions: a helix-bundle rock along with movement of TM1a, 1b, 5a, 6a, and 10a. (F) Rescaled PCA weights for all six hb-DDMs for LeuT, including the three one-step transitions (left), the two possible two-step transitions, and the three-step transition from O_op_ to I_op_. Weights are colored according to their color in panel (E).

### Six motions are commonly observed in LeuT-fold transporters

Based on visual inspection, we hypothesized that these complex hb-DDMs are themselves combinations of different “building block” motions. To identify these building blocks, we performed sparse principal component analysis (PCA), which reveals six simple motions that explain much of the variability among hb-DDMs, with a mean squared reconstruction error of 0.90 Å (Figure 3D). Each motion corresponds either to swinging of one of five half-helices—1a, 1b, 5a, 6a, or 10a—or to an intracellular bundle-hash rocking motion of 1a, 2, 6b, and 7 against 4, 5a, 8 and 9 (Figure 3E). Each individual hb-DDM can then be expressed as a weighted sum of these building block motions, as is illustrated for the six possible pairwise comparisons of the four LeuT conformations (Figure 3F). Motions of TM1b (green), TM6a (orange), and TM10a (red) participate in outward-closing (O_op_→O_oc_), while the O_oc_→I_oc_ transition primarily features the hash-bundle rock (gray) and the swinging of TM5a (yellow). By contrast, the inward-opening motion (I_oc_→I_op_) is dominated by the swinging motion of TM1a (blue). The weights for all 22 pairwise structural comparisons are in Figure S7. Below we use these sparse PCA weights alongside the three types of DDMs to address whether conformational motions in the LeuT-fold superfamily are similar across homologs.

### Bundle-hash rock is common to all outward-to-inward transitions

Seven transporters have an O_op_→I_op_ comparison available: LeuT, SERT, BetP, Mhp1, MntH, KCC1, and LAT1. In all these proteins, the sparse PCA weights indicate that the outward-to-inward transition involves a substantial hash-bundle rock motion; this is the largest common PCA weight contribution in these comparisons (Figures 4A and S7). The rock PC involves bundle helices TM1a, 2, 6b, and 7 moving relative to scaffold helices (TM4, 8, and 9 of the hash, and 5a and 10b of the arms; Figure 3B), and it features prominently in some hb-DDMs like that of the Mhp1 O_oc_→I_op_ transition (Figure 4B), while it is present but less striking in others, like that of the BetP O_oc_→I_op_ transition (Figure 4C). The rock motion contributes to closing of the outer vestibule and leads to a separation of the bundle and hash to open the intracellular vestibule. This is readily apparent in a superposition of the O_oc_ and I_op_ conformations of Mhp1, in which this rock motion leads to a ∼20° rotation of the bundle relative to the hash (Figure 4B). The superposition of Mhp1 structures suggests that the fulcrum of rotation is in the outer leaflet of the membrane, leading to larger distance differences between the inner bundle and hash. This is consistent with the fact that all half-helices involved in the rock PC are in the inner leaflet of the membrane (Figures 4A and 4C) and suggests that the synchronized rock of the bundle is biased toward the inner half of the proteins. Although the bundle-hash rock of BetP has been described as small relative to other LeuT-fold transporters (Supplementary note; ^11^), the ∼5° rock motion (Figure 4D) contributes the most of any PC to the BetP O_oc_→I_op_ transition (Figures 4A and 4E).

**Figure 4.**
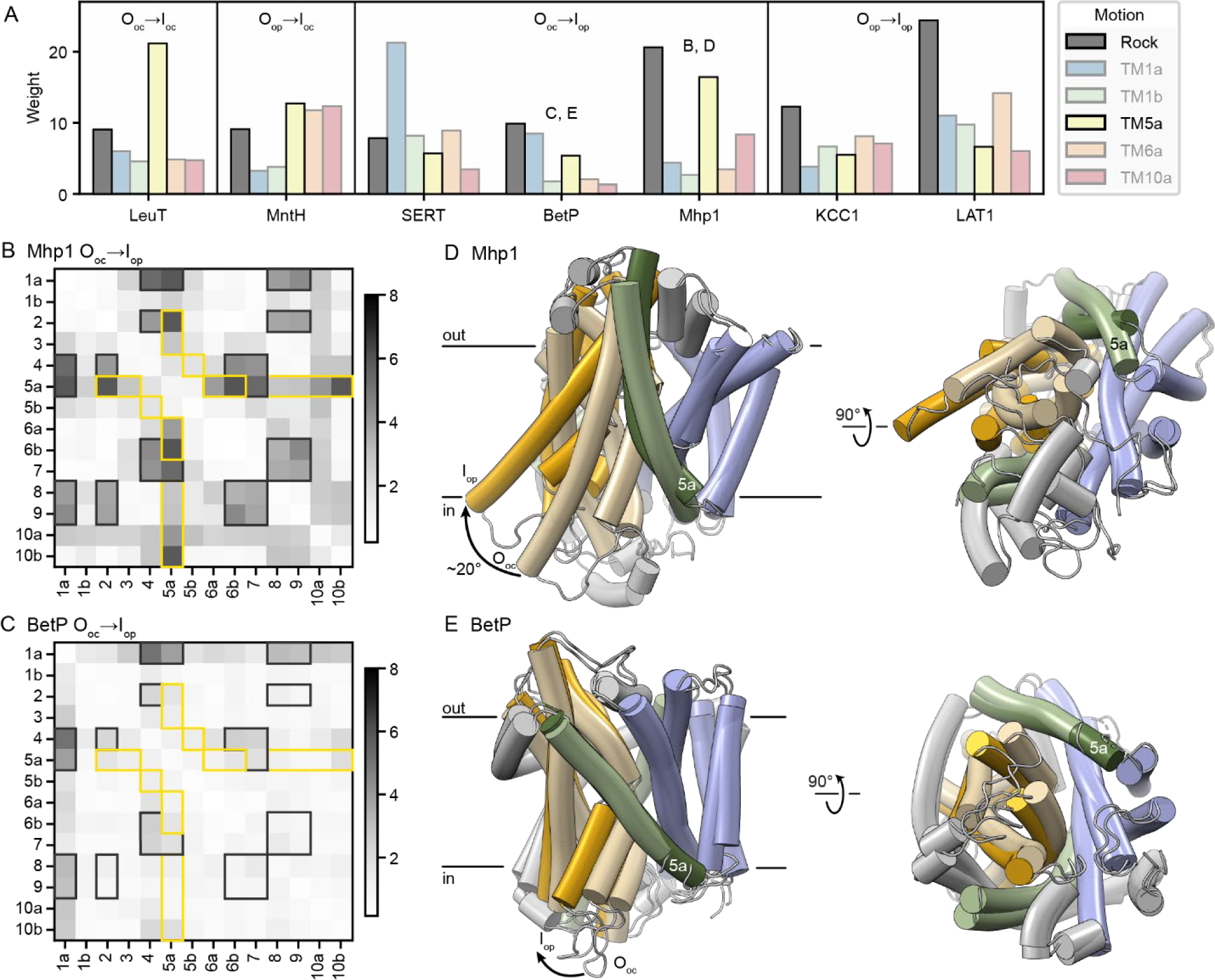
Rock motion of the bundle relative to the hash is a common feature of the outward-to-inward transition in LeuT-fold transporters. (A) Sparse PCA weights for a set of outward-to-inward transitions: the one example of an O_oc_→I_oc_ transition (LeuT), and transitions for other proteins that incorporate the O_oc_→I_oc_ motion: O_op_→I_oc_ for MntH, O_oc_→I_op_ for SERT, BetP, and Mhp1, and O_op_→I_op_ for KCC1 and LAT1. In each case, the helix-bundle rock (grey) and TM5a (yellow) PCs are highlighted. The letters above the histograms refer to the respective figure panels. (B, C) hb-DDMs for the O_oc_→I_op_ transitions of Mhp1 (B) and BetP (C), with the elements contributing to the helix-bundle rock and TM5a weights highlighted in dark grey and yellow, respectively. (D) Cartoon representations of the O_oc_ and I_op_ structures of Mhp1, which were superimposed using their hash helices (RMSD = 0.9 Å over 97 Cα atoms). The Mhp1 bundle rotates ∼20° to open the intracellular vestibule, essentially as a rigid body (RMSD = 0.6° over 110 Cα atoms). The I_op_ bundle helices are in deeper yellow and TM5 in deeper green to highlight the movements. On the left, the approximate position of the membrane for the O_oc_ structure is indicated (determined using the OPM database^18^). The illustration on the right is rotated 90°, viewing the proteins from the cytosolic side. (E) Cartoon representations of the O_oc_ and I_op_ structures of BetP, which were superimposed using their hash helices (omitting TM8 based on the clustering analysis in Figure S7; RMSD = 1.0 Å over 60 Cα atoms). The BetP bundle rotates ∼5° to open the intracellular vestibule, with an RMSD of 1.0° over 110 Cα atoms when the bundles are superimposed.

The outward-to-inward transition also includes a shift of TM5a away from the bundle to open the intracellular vestibule. This is particularly prominent in LeuT, Mhp1, and MntH (Figures 4 and S7). The TM5a PC is generally most prominent in the O_oc_→I_oc_ and I_oc_→I_op_ transitions. SERT is an exception, as its TM5a moves earlier in the transport cycle, constituting the most prominent weight of the O_op_→O_oc_ transition.

### Only some LeuT-fold transporters show a large TM1a swing to open inward

While the outward-to-inward transition is consistent for all transporters we analyzed, the motions that precede and follow that step vary. In several transporters, TM1a swings upward during the transition to I_op_, opening the intracellular vestibule to provide access to the substrate-binding site from the cytosol (Figure 5 and Table S1). The TM1a PC makes up by far the greatest weight for the LeuT I_oc_→I_op_, MntH I_oc_→I_op_, and SERT O_oc_→I_op_ transitions and is also substantial for LAT1 (O_op_→I_op_) and BetP (O_oc_→I_op_; Figure 5A). For the transporters without I_oc_ structures, this TM1a swing likely still dominates the I_oc_→I_op_ transition, regardless of whether the I_oc_ state is a stable intermediate. The corresponding hb-DDMs show that TM1a moves relative to most LeuT-fold helices (Figures 5B-C and S3). The ho-DDMs confirm the upward swing motion: the LeuT, MntH, SERT, BetP, and LAT1 TM1a helices move away from most helices—especially the inner halves of the hash helices—and toward TM5b and the outer halves of TM7 and TM8 (Figure S3). By contrast, TM1a moves little in the transition to I_op_ of SGLT, Mhp1, or KCC1, as they have low weights for the TM1a PC (Figure 5A) and the hb-DDMs show a relatively static TM1a, as illustrated for SGLT in Figure 5D.

**Figure 5.**
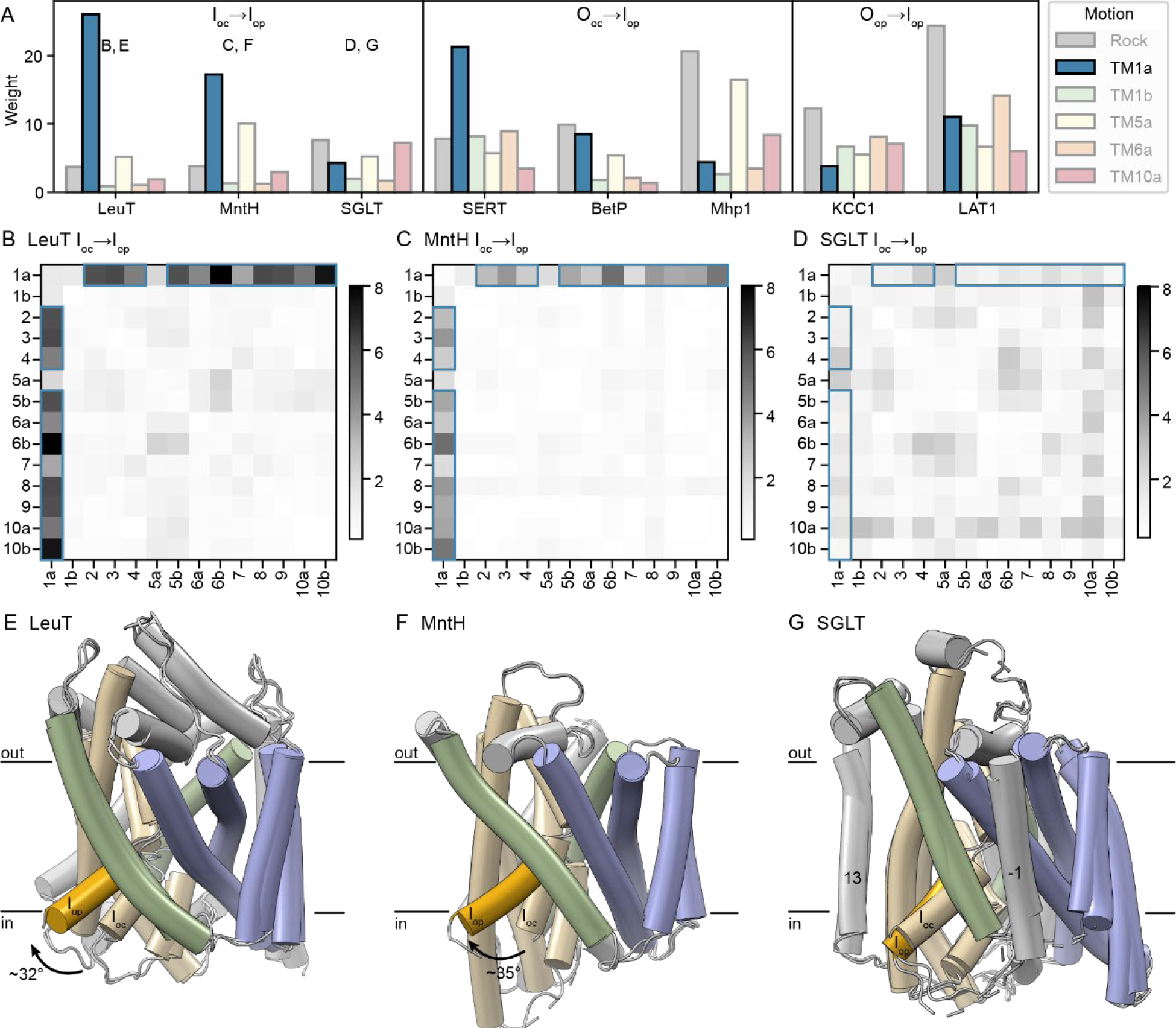
Many but not all LeuT-fold transporters close the intracellular vestibule primarily through a swing of TM1a. (A) Sparse PCA weights for a set of transitions that open the intracellular vestibule, showing the I_oc_→I_op_ transition for the three proteins with both such structures, or the O_oc_→I_op_ or O_op_→I_op_ transition for proteins with no available I_oc_ (and O_oc_) structures. In each case, the PC representing the TM1a motion is highlighted and shown in blue. The letters above the histograms refer to the respective figure panels. (B-D) hb-DDMs for the I_oc_→I_op_ transition for LeuT (B), MntH (C), and SGLT (D), with the RMSDDs most associated with the TM1a PC highlighted in blue. (E-G) Cartoon representations of the I_oc_ and I_op_ conformations superimposed using their hash helices for LeuT (E; RMSD = 0.5 Å over 107 Cα atoms), MntH (F; RMSD = 0.3 Å over 93 Cα atoms), and SGLT (G; RMSD = 0.8 Å over 103 Cα atoms). TM1a in the I_op_ conformation is in deeper yellow to highlight its upward swing. The TM1a of LeuT swings by ∼32°, that of MntH by ∼35°, whereas the SGLT TM1a does not substantially move. The extra helices in SGLT that likely impinge on TM1a motion—TM-1 and TM13—are labeled.

The I_oc_ and I_op_ structures of LeuT and MntH superimposed on their hash exemplify the isolated TM1a motion, with >30° swings in both cases (Figure 5E-F), whereas in SGLT TM1a barely moves (Figure 5G). Of note, the nearby position of some of the extra helices in SGLT (TM-1 and TM13) may limit the range of motion of TM1a (Figure 5G). Instead, the SGLT I_oc_→I_op_ comparison shows the outer bundle moving closer to the hash, and the inner half of TM8 moving away from the rest of the protein to open the intracellular vestibule (Figures 5G and S3J). This suggests that the swinging of TM1a is a common although not requisite path to inward opening in the LeuT-fold.

### TM1b, TM6a, and TM10a can move independently to close the outer vestibule

Based on the DDMs, outward occlusion can occur through movement of any of TMs 1b, 6a or 10a into the outer vestibule (Figure 6). In the O_op_→O_oc_ transitions of Mhp1, BetP, and AdiC, arm helix TM10a moves toward the outer bundle helices (Figures 6A, 6B, 6D, 6E, and 6G). This motion is isolated to TM10a and does not include TM10b, as seen clearly in the O_op_→O_oc_ hb-DDM of Mhp1 (Figure 6E), where the motion of TM10a alone closes the outer vestibule. However, TM10a does not move in LeuT and SERT (Figures 6A, 6C, and 6F), showing that TM10a movement is not necessary for outward occlusion of the LeuT fold.

**Figure 6.**
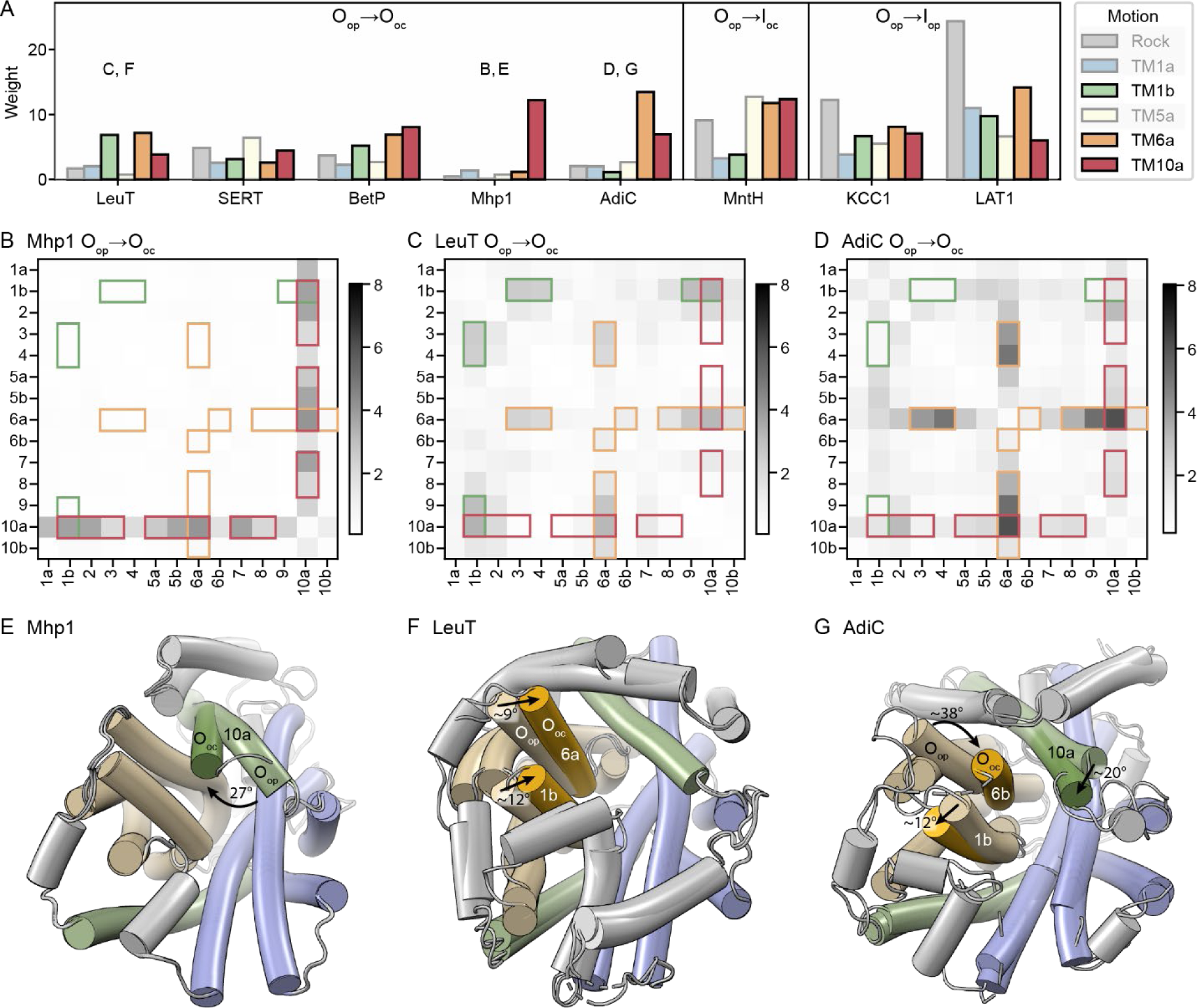
Different combinations of motions in TM1b, TM6a, and TM10a close the outer vestibule. (A) Sparse PCA weights for a set of transitions that close the outer vestibule, showing the O_op_→O_oc_ transition for the five proteins with both such structures, or the O_op_→I_oc_ or O_op_→I_op_ transition for proteins with no available O_oc_ (and I_oc_) structures. In each case, the PCs representing the TM1b, TM6a, and TM10a motions are highlighted and shown in green, orange, and red, respectively. The letters above the histograms refer to the respective figure panels. (B-D) hb-DDMs for the O_op_→O_oc_ transition for Mhp1 (B), LeuT (C), and AdiC (D), with the RMSDDs most associated with the TM1b, TM6a, and TM10a PCs highlighted in green, orange, and red, respectively. (E-G) Cartoon representations of the O_op_ and O_oc_ conformations superimposed using their hash helices, for Mhp1 (E; RMSD = 0.2 Å over 97 Cα atoms), LeuT (F; RMSD = 0.4 Å over 107 Cα atoms), and AdiC (G; RMSD = 0.8 Å over 89 Cα atoms). For the cases where they move substantially, TM1b and TM6a in the O_oc_ conformation are in deeper yellow, and TM10a is in deeper green. The approximate angular motions are indicated and marked by arrows.

The O_op_→O_oc_ transition can also include movements of two outer bundle helices, TM1b and TM6a, toward the outer hash. In LeuT, SERT, and BetP, TM1b moves closer to the outer scaffold (Figures S3A-S3D). This motion is also seen in the O_op_→I_oc_ transition of Nramp and the O_op_→I_op_ transitions of KCC1 and LAT1, for which there are no O_oc_ structures (Figures 6A and S3F-S3H). However, TM1b does not move in the Mhp1 O_op_→O_oc_ transition (Figures 6B and 6E), and TM1b moves away from the hash rather than towards it in AdiC (Figures 6D and 6G). TM6a also moves closer to the hash in all O_op_→O_oc_ transitions except for SERT and Mhp1. In SERT, whose O_op_→O_oc_ transition shows limited motions (Figure S3C), TM6a moves later in the O_oc_→I_op_ transition (Figure S7). This TM6a motion therefore often but not always accompanies TM1b motion, and it can also be paired with TM10a motion, highlighting how different proteins employ different combinations of TM1b, TM6a, and TM10a motions to close the outer vestibule. Overall, this mechanistic diversity suggests that there is no one way that LeuT-fold transporters close the outer vestibule to occlude the substrate-binding site. Rather, the flexibility of the fold allows for differing usage of the TMs 1b, 6a, and 10a, as reflected by the fact that motion of each of these helices is represented by a different PC.

### Either the bundle or hash or both can rotate relative to the membrane

We determined above that the bundle and hash rock relative to one another in LeuT-fold transporters. What remains unclear is which structural elements move and which are static relative to the membrane, which represents a natural fixed reference plane. To determine whether the bundle or hash move relative to the membrane in the O_op_→I_op_ transition, we first used the Orientations of Proteins in Membranes (OPM) database and Positioning of Proteins in Membranes (PPM) server^18^ to predict the membrane position for the O_op_ and I_op_ structures of each protein with both conformations available. We then superimposed the two structures using either the hash or bundle helices and determined the resulting angle, θ, between the membrane planes (Figures 7A-7C and Table S7). The direction of rotation is generally consistent with both the bundle and hash rotating up and into the outer vestibule in the O_op_→I_op_ transition. The only exception is the small 4° rotation when superimposing according to the SERT hash, which is consistent with the hash rotating into the intracellular vestibule.

**Figure 7.**
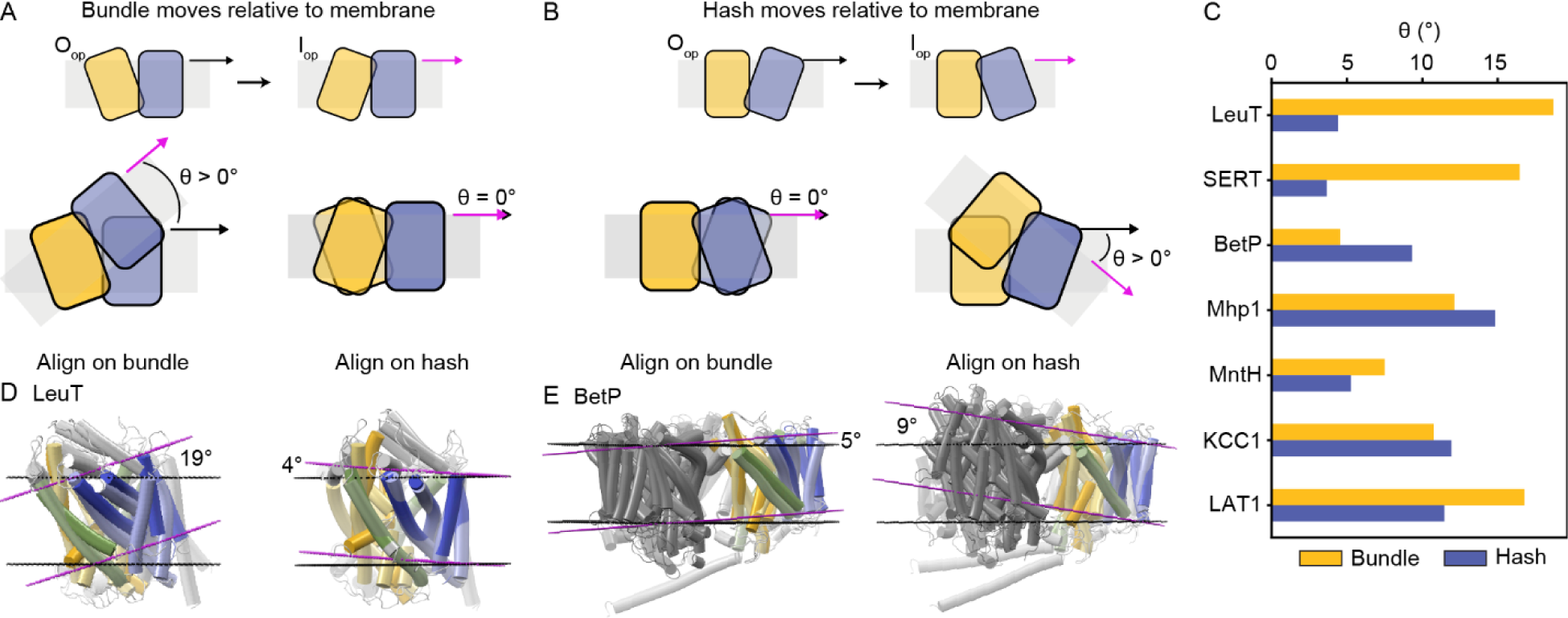
Different LeuT-fold transporters have the bundle or hash rotate more relative to the membrane. (A) Schematics of alternating access enabled by the bundle (yellow) moving while the hash (blue) is fixed relative to the membrane (grey). The corresponding O_op_ and I_op_ conformations are illustrated at the top. At the bottom, the two conformations are superimposed using their bundle (left) which results in a rotation of the corresponding membrane planes by the angle θ, measured between the black (O_op_) and magenta (I_op_) vectors, or hash (right) which leaves the corresponding membranes static. (B) Schematics of alternating access enabled by the hash moving while the bundle remains fixed relative to the membrane. The O_op_ and I_op_ conformations are at the top. At the bottom, the two conformations are superimposed using their bundle (left) which leaves the corresponding membranes static, or hash (right) which results in a rotation of the corresponding membrane planes. (C) Measurements of the angle (θ) between the two predicted membrane planes from bundle-based (yellow) or hash-based (blue) superpositions of the O_op_ and I_op_ structures of different LeuT-fold transporters. (D) Bundle-based (left) or hash-based (right) superpositions of the O_op_ (black membrane planes) and I_op_ (magenta membrane planes) structures of LeuT, showing that alternating access in LeuT arises primarily from the bundle moving relative to the membrane. (E) Bundle-based (left) or hash-based (right) superpositions of the O_op_ and I_op_ structures of BetP, showing that alternating access in BetP arises primarily from the hash moving relative to the membrane (color-coded as in panel D). The whole BetP trimer is shown, with the protomer used in the superpositions colored and the other two protomers in medium grey.

The magnitude of rotation of each structural element differs substantially between the different transporters. The greater θ is for the superposition based on a given structural element, the more that element rotates relative to the predicted membrane plane in the O_op_→I_op_ transition. For LeuT and SERT, the bundle rotates more relative to the membrane (Figures 7A and 7C). For BetP, the hash rotates more—probably because the bundle is constrained within the trimer interface (Figures 7B and 7D). For Mhp1, KCC1, and LAT1, both rotate substantially, and for MntH both rotate modestly. Only for LeuT, SERT, and BetP is there a greater than 50% difference between the values of θ for the bundle and the hash. This indicates that the LeuT-fold conformational mechanism commonly accommodates movement relative to the membrane plane of both the bundle and the hash during the outward-to-inward transition and that which structural element moves most relative to the membrane varies considerably among superfamily members.

## DISCUSSION

To determine the extent to which the variation of conformational movements is accommodated within the LeuT fold, we have adapted a superposition-position free approach to analyzing structural differences—the distance difference matrix (DDM)—to compare the transport cycles of nine LeuT-fold transporters with structures available for more than one conformation. By binning the data according to α-helices to enable comparisons across distantly related proteins and decomposing the resulting set of 22 hb-DDMs into sparse PCs, we identified six recurring conformational movements as the transporters transition from O_op_ to I_op_: rocking of the bundle and hash relative to one another to achieve an inward-facing state, TMs 1b, 6a, and 10a each moving into the outer vestibule independently, TM5a moving away from the intracellular vestibule, and TM1a swinging up and away from the intracellular vestibule. The well-established rocking-bundle motion occurs in all transporters, so this motion is likely required to transition from an outward orientation to an inward one. However, in different transporters either the bundle or the hash can be more mobile relative to the membrane plane. The opening or closing of each vestibule is governed by motions of multiple individual helices—TMs 1b, 6a, and 10a for the outer vestibule and TM1a and TM5a for the intracellular vestibule—with each helix contributing to different degrees in different transporters. This redundancy could enable each protein to sample from a menu of conformational movements to achieve a combination most advantageous for their environment and the transport of their specific substrate.

### The bundle-hash rock is conserved in the LeuT fold

There is no consensus on the extent to which the overall conformational change mechanism of the LeuT-fold superfamily is conserved. A previous quantitative analysis found that the relative arrangement of the TMs is more tightly associated to the conformational state than to the particularities of each protein^3^, but deviations from the canonical rocking-bundle model for BetP^11^ and KCC1^13^ have cast doubt on its universality. A recent review asserted that the contribution of the bundle to alternating access is “far from universal”^10^. Individual helix movements that vary from protein to protein were identified as facilitating alternating access in most cases, suggesting that “divergence is the rule”^10^.

Our systematic analysis shows that the core bundle-hash rock motion is conserved as a dominant feature of the outward-to-inward transition across all LeuT-fold superfamily members, even BetP and KCC1, although BetP does indeed have the smallest rotation between the hash and bundle. The confusion about whether the bundle-hash rock is an obligate element of the LeuT-fold transport cycle may stem from two facts that our analysis underscores: (i) the bundle is often not a rigid body as several bundle helices (TMs 1a, 1b, and 6a) move independently in addition to the global rock motion, and (ii) LeuT-fold transporters exhibit substantial variation in whether the bundle, the hash, or both rotate relative to the *membrane* (not whether they rotate relative to each other). Initial analyses of the outward- and inward-facing conformations for both BetP^11^ and KCC1^13^ suggested that those proteins may have a fundamentally different mechanism to achieve alternating access due to a perceived greater motion of the hash rather than the bundle. However, this observation is dependent on the reference frame. Our reference-free quantitative analysis of the hb-DDMs categorizes the movements of the hash seen in BetP and KCC1 in the same PC as the rocking bundle observed in LeuT and others. This indicates that for all LeuT-fold transporters studied, the bundle and hash must move relative to one another in a similar manner in the outward-to-inward transition. When considering the membrane as the reference plane, BetP and KCC1 do indeed have a more mobile hash than LeuT, although this is also true of Mhp1 and LAT1. Therefore, the bundle-hash rock seems to be an obligate motion of the LeuT-fold transport cycle, but the direction of this motion relative to the membrane differs among superfamily members.

The ability for either the bundle or the hash to move relative to the membrane as part of the rocking motion may serve as a functional adaptation. BetP, whose hash moves more than the bundle, has a network of molecular interactions linked to its trimeric state that restricts the flexibility of bundle and regulates transport, an adaptation that enables it to dynamically respond to environmental changes as part of its role in osmosensing and osmoregulation^19,20^. The KCC1 hash also moves substantially relative to the membrane (Figure 7C) due to a KCC-specific disulfide bond between TM2 of the bundle and TM11, a helix outside of the LeuT-fold core^13^. The inhibitor-bound O_op_ structure of KCC1 adopts an alternate dimeric state that relies on a different interface. The dimer status has been hypothesized to be an adaptation that enables regulation of KCC1 by kinases and other cellular factors^13^.

### LeuT-fold transporters sample from a set of common individual helix motions to generate mechanistic diversity

While divergence may not be the rule for the rocking bundle, it is for the opening and closing of the vestibules at each end of the transport cycle. We find that the transporters close the extracellular vestibule through different combinations of the movement of individual helices TMs 1b, 6a, and 10a, and close the intracellular vestibule either with or without a large swing of TM1a. Motions of all these helices have been previously noted, but often with somewhat different conclusions. Previous work has indicated that TM1b and TM6a move to close the outer vestibule^10,21,22^, but our analysis disputes the idea that movement of these helices is necessarily coupled. Our results also suggest that these motions may be more common than previously thought; the movements of TM1b and TM6a were not reported for KCC1^13^, but these motions contribute substantially to the KCC1 O_op_→O_oc_ hb-DDM. Also in this transition, TM10a swings away from the hash and toward the bundle to fill the outer vestibule in AdiC, BetP, KCC1, LAT1, Mhp1, and MntH—all but the NSS transporters LeuT and SERT^3,10,17,23^.

Our analysis identifies movement of TM5a away from the inner vestibule as a major contributor, along with the bundle-hash rock, to the outward-to-inward transitions for all seven proteins with such comparisons available. These results are consistent with an understanding in the field that TM5 movement away from inner portion of the bundle is broadly important to open the intracellular vestibule^3,10,17,21,24^. While TM5a moves in all transporters during outward-to-inward transitions, the movement is greater for certain transporters like LeuT and Mhp1 (Figure 4). MD simulations and mutational analysis with LeuT indicate that the presence of substrate in the outer vestibule transduces a signal through the transporter which promotes TM5 unbending^8^. The greater magnitude TM5 movement in LeuT could facilitate this functional adaptation.

Likewise, previous studies have highlighted the importance of TM1a movement to inward opening, but doubt has existed about its physiological relevance because distance distributions measured by double electron-electron resonance spectroscopy in LeuT were inconsistent with a large displacement^22^. Our analyses indicate that the TM1a swing is a major contribution in the I_oc_→I_op_ transition for the proteins that exhibit this movement, LeuT, SERT, LAT1, and MntH. The movement has been observed in I_op_ structures determined both in detergent (LeuT, SERT, LAT1) and in lipid bilayers (MntH), and using both X-ray crystallography (LeuT, MntH) and cryogenic electron microscopy (SERT, LAT1). Substantial TM1a flexibility was also observed in LeuT using single molecule-Fluorescence Resonance Energy Transfer^8^. The TM1a swing is also vital for Co^2+^ transport in DraNramp^25^, likely facilitating substrate release. The diversity of circumstances in which this movement has been observed suggests that it is not an artifact and is indeed an important conformational change for certain LeuT-fold proteins. However, multiple members of the LeuT-fold superfamily have alternate mechanisms of inward-opening, suggesting that this mechanism is not universal. For instance, non-LeuT-fold TMs −1 and 13 of SGLT may limit TM1a swinging (Figure 5F). TM13 of the human homologs SGLT1 and SGLT2 binds to MAP17, an essential subunit for hSGLT2 glucose uptake activity that promotes cell surface expression^26^.

There are no major repeatedly observed motions outside of the ones captured by the top six PCs in our analysis. In AdiC, TM2 moves (along with and behind TM6a) toward the outer vestibule in the O_op_→O_oc_ transition^23^. This is borne out in the corresponding DDMs although no other analyzed transporter clearly displays individual TM2 movement in their DDMs (Figure S3), and no functional studies or structures support the existence of this movement in other LeuT-fold transporters^10^. Our DDM-based analyses show no important motion for TM7. However, electron paramagnetic resonance and hydrogen-deuterium exchange/mass spectrometry experiments have suggested that TM7 may partially unfold in the LeuT I_op_ conformation, a finding contrary to the experimental structures^10,22^. Further experiments will be necessary to determine whether this potential movement is physiologically relevant and whether it occurs in other transporters.

In sum, in addition to variation in the magnitude and orientation relative to the membrane of bundle-hash rocking, each LeuT-fold transporter samples from a set of common motions to produce an overall transport cycle that accommodates the constraints of its substrate selectivity and the membrane environment.

### Comparing binned DDMs across entire protein families is applicable to a diverse range of analyses

We can make the above conclusions about which parts of each protein move relative to one another by using superposition-free structural comparisons with hb-DDMs. While such methods have been used previously on a small scale^17^, we present the first systematic use of hb-DDMs to analyze motions across a protein superfamily, an approach that is applicable to any protein family that changes conformations within a core structural scaffold. The choice of binning based on helices is particularly appropriate for helical membrane proteins, but the framework is easily adapted to bin and compare based on other secondary structure elements or even entire domains for large multi-domain proteins. Angle difference matrices could also be generated using a similar computational approach, which would focus analyses on rotational movements of structural elements. Furthermore, while DDMs require structures in multiple conformations, distance matrices (DMs) require only a single conformation. The LeuT-fold transporters with structures in only one conformation (which were excluded from our analyses) could be included in an analysis of DMs to compare a given conformation between different proteins.

Our DDM workflow, or adaptations like the ones suggested above, could also be integrated with other computational techniques. For example, the different conformational states could be determined through molecular dynamics (MD) simulations, or through predictions like those of AlphaFold^27^. While new methods have facilitated prediction of multiple conformation states of proteins via AlphaFold^28,29^, it remains to be seen whether the transitions between these predicted states are biologically relevant. As we have demonstrated, our workflow showed in a systematic manner that LeuT-fold transporters exhibit differences in their conformational movements that relate to their varied functions and demonstrates the applicability of binned DDMs to analyze motions in protein families.

## Supporting information

Supplementary Data File 1. Annotated list of structures.

## ACKNOWLEDGEMENTS

We acknowledge members of the Gaudet Lab for useful discussions and Casey Zhang for providing some of the initial python scripts. This work was funded in part by NIH grant R01GM120996 (R.G.), the NSF-Simons Center for Mathematical and Statistical Analysis of Biology at Harvard (award number 1764269) and the Harvard Quantitative Biology Initiative (S.P.B.), and the Harvard College Research Program (J.A.L.).

## AUTHOR CONTRIBUTIONS

R.G. supervised the work and provided resources. J.A.L., M.A.G. and R.G. conceived and designed the project, and assembled and curated the structure dataset. J.A.L. performed the RMSD analyses and the membrane orientation alignments. S.P.B. performed the statistical and clustering analyses. J.A.L. and S.P.B. developed the code and calculated the DDMs. J.A.L., S.P.B., and R.G. prepared all the data presentation and visualization and wrote the manuscript. All authors reviewed and edited the full manuscript.

## STAR METHODS

### RESOURCE AVAILABILITY

#### Lead contact

- Further information and requests for resources should be directed to and will be fulfilled by the lead contact, Rachelle Gaudet.

#### Materials availability

- Materials used in this study are publicly available at the Protein Data Bank.

#### Data and code availability

- All original code has been deposited on Github at https://github.com/GaudetLab/pyDDM.
- Any additional information required to reanalyze the data reported in this work paper is available from the lead contact upon request.

## METHOD DETAILS

### Compilation of LeuT-fold structures

We created a comprehensive list of published LeuT-fold structures by combining the results of DALI^12^ searches using each of the 12 structures listed in Table S5. Each search yielded between 238-298 high-scoring hits (readily identified from a large drop in both the Z-score and the “lali” value of number of aligned residues for the next-best scoring hit), with three searches having the maximal value of 298 hits, and the combined non-redundant list including these same 298 hits, providing confidence that all LeuT-fold structures available on that date (June 11, 2021) were captured by these searches. The annotated list, including eight added MntH structures and one added KCC1 structure, is in Supplementary Data File 1.

The 298 hits correspond to 207 unique PDB ID codes (some structures have multiple chains, which we treat here as individual structures). Using UniProtKB and manual editing, we mapped these structures to 36 unique genes of 28 different proteins (for several proteins, structures of more than one orthologs are available), as summarized in Table 3.

### Conformation assignment of the LeuT-fold structures

For each transporter with multiple structures, we used the cealign function in PyMOL with default parameters to calculate the average Cα RMSD for each pair of structures, yielding an RMSD matrix. Each matrix was clustered using k-means, and the clusters were annotated and re-ordered according to transporter conformation. For each cluster, we used the literature to identify the conformation of at least two of the structures, then annotated all structures of that cluster accordingly if the RMSD between structures in the cluster was below 0.5. A few conformations were labeled as “closed” or “occluded” in the literature, but visual inspection suggested they were closest to O_oc_ in our classification scheme. In general, these structures had RMSD values of at most 2 Å with respect to O_oc_ structures and clustered with O_oc_ structures. Even when we increased the number of clusters, the O_oc_ and “closed” or “occluded” structures did not cluster as separate groups but instead split into groups that contained both O_oc_ and “closed” or “occluded.” Thus, we categorized these “occluded” and “closed” structures as O_oc_.

For each protein with structures in multiple conformations, one structure in each conformation was chosen as representative based on the following priorities: (1) choose structures derived from the same organism; (2) avoid structures with an atypical RMSD relative to other structures grouped in the same conformation; (3) choose the structure with the highest resolution; (4) structures with a disordered TM in the LeuT-fold core were replaced with the next best structure of the same conformation. The chosen structures are listed in Table 3.

### Distance-difference matrices

We generated DDMs comparing each possible conformation pair for each protein by first computing all the pairwise Cα-Cα distances (*d_ij_*) between all residues between the two structures. For consistency in our comparisons, we considered the half conformational cycle starting from O_op_, moving through O_oc_ then I_oc_, and ending at I_op_, and assumed that these conformations are not substantially altered by the presence or absence of substrate at the resolution of our analysis (i.e., we ignored information about presence of substrate). All DDMs were calculated as *d_ij_ ^B^*-*d_ij_ ^A^* for an A→B comparison.

To generate ho-DDMs and hb-DDMs, we needed to select residue ranges for the LeuT-fold core TM helices for each protein (Table S4). We used PROMALS3D^33^, to generate a rough structure-based sequence alignment, and used the alignment used to set uniform residue cutoffs for each helix as a starting point. We then manually adjusted these ranges by visual inspection of the structures and only including residues that are helical in all available conformations of the respective protein. The first and last helix of each repeat (TM1, 5, 6, 10) were split for each structure. TM1 and TM6 contain breaks in all LeuT-fold transporters. For TM5 and TM10, only some transporters contain breaks in those helices; we structurally aligned all transporters and chose the residue best aligned with the breaks as the break position for other transporters. The N- and C-terminal halves of the split helices are labeled “a” and “b,” respectively. TM helices were numbered relative to their position within the LeuT-fold core, regardless of the presence of additional N- or C-terminal TM helices in some transporters. ho-DDMs display only the defined TM residues, with inserted black vertical and horizontal lines marking the start and end of each helix.

To make hb-DDMs, we adapted a published Python script^17^ that takes the norm of all distance differences between two helices ℎ_1_ and ℎ_2_:

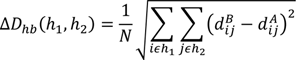

where *N* is the product of the lengths of ℎ_1_ and ℎ_2_ and 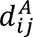 is the distance between residues *i* and *j* in structure *A*. We refer to the values in hb-DDMs as root mean squared distance differences, or RMSDDs.

### Rigid body identification via helix clustering

To identify rigid bodies relative to a given transition, we performed hierarchical clustering on the helices with the hb-DDM as an input distance matrix using scipy’s “hierarchy” module^34^. We first converted the hb-DDM to a proper distance matrix between helices by setting the diagonal entries to zero and then used the “average” (UPGMA) method in scipy’s hierarchical clustering module to define a linkage matrix, from which we visualized clustered hb-DDMs as clustermaps in seaborn^35^.

### Sparse principal component analysis (PCA)

Each of the 22 14×14 hb-DDMs was flattened into a 196-feature vector and sparse PCA was performed on these vectors with sklearn (v1.2.1)^36^, where the reconstruction loss of the data projection is balanced with an L1 norm on the components. We set a sparsity parameter of α=2 and used the least angle regression method to solve the lasso problem. We set the number of components to six, although we also generated components for a range of other values of *n*. The resulting components were all strictly positive or strictly negative; for subsequent analyses, all were fixed to be strictly positive.

Purely for interpretability, the PCA projections of each original flattened DDM were shifted by a constant vector corresponding to the projection of a matrix of all zeroes. This redefines the origin in the PCA latent space to correspond to a lack of motion such that the projections are all nonnegative and defined absolutely rather than relative to the mean of the dataset. We also used non-negative matrix factorization to express the matrices as purely positive sums of the top PCs (fixing H to be the top 6 sparse PCs with sklearn’s “custom” protocol) and this gave nearly the exact same result; all figures show the “rescaled” PCA projections, but the results could be equivalently thought of as the nonnegative regression coefficients on the components. Using non-negative matrix factorization to infer the transformation matrix H, however, did not lead to interpretable results, even setting a strict L1 norm to induce sparsity.

### Membrane orientation alignments

We downloaded each O_op_ and I_op_ structure with its predicted membrane orientation from the OPM server^18^. If the native oligomerization state (determined by previous findings) differed from the available structures, or if the OPM server contained a curved membrane, we used PPM 3.0 with default parameters to predict the membrane orientation for the native oligomer. We aligned the O_op_ and I_op_ structures with their predicted membranes using the pair_fit command according to either the bundle, hash, or core helices. To determine the resulting angle, *θ*, between the predicted membrane planes, we first calculated the normal vectors, *v*, for the membrane planes using the *x,y,z* coordinates of three dummy atoms from the intracellular membrane plane:

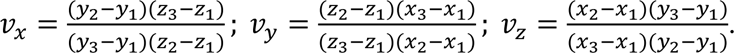

We then calculated *θ* as the angle between the normal vectors for the O_op_ and I_op_ structures, *v_O_* and *v_I_*, respectively:

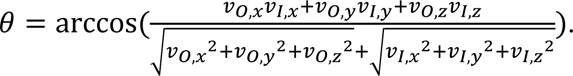

## QUANTIFICATION AND STATISTICAL ANALYSIS

Details of the quantification and statistical analysis can be found in the figure and table legends and the corresponding STAR methods sections.

## References

1. Vastermark, A., Wollwage, S., Houle, M.E., Rio, R., and Saier, M.H., Jr. (2014). Expansion of the APC superfamily of secondary carriers. Proteins 82, 2797–2811. 10.1002/prot.24643.

2. Koh, P.O., Undie, A.S., Kabbani, N., Levenson, R., Goldman-Rakic, P.S., and Lidow, M.S. (2003). Up-regulation of neuronal calcium sensor-1 (NCS-1) in the prefrontal cortex of schizophrenic and bipolar patients. Proc Natl Acad Sci U S A 100, 313–317. 10.1073/pnas.232693499.

3. Jeschke, G. (2013). A comparative study of structures and structural transitions of secondary transporters with the LeuT fold. Eur Biophys J 42, 181–197. 10.1007/s00249-012-0802-z.

4. Yamashita, A., Singh, S.K., Kawate, T., Jin, Y., and Gouaux, E. (2005). Crystal structure of a bacterial homologue of Na^+^/Cl^-^-dependent neurotransmitter transporters. Nature 437, 215–223. 10.1038/nature03978.

5. Forrest, L.R., Kramer, R., and Ziegler, C. (2011). The structural basis of secondary active transport mechanisms. Biochim Biophys Acta 1807, 167–188. 10.1016/j.bbabio.2010.10.014.

6. Edwards, N., Anderson, C.M.H., Conlon, N.J., Watson, A.K., Hall, R.J., Cheek, T.R., Embley, T.M., and Thwaites, D.T. (2018). Resculpting the binding pocket of APC superfamily LeuT-fold amino acid transporters. Cell Mol Life Sci 75, 921–938. 10.1007/s00018-017-2677-8.

7. Forrest, L.R., and Rudnick, G. (2009). The rocking bundle: a mechanism for ion-coupled solute flux by symmetrical transporters. Physiology (Bethesda) 24, 377–386. 10.1152/physiol.00030.2009.

8. Zhao, Y., Terry, D., Shi, L., Weinstein, H., Blanchard, S.C., and Javitch, J.A. (2010). Single-molecule dynamics of gating in a neurotransmitter transporter homologue. Nature 465, 188–193. 10.1038/nature09057.

9. Koldso, H., Noer, P., Grouleff, J., Autzen, H.E., Sinning, S., and Schiott, B. (2011). Unbiased simulations reveal the inward-facing conformation of the human serotonin transporter and Na^+^ ion release. PLoS Comput Biol 7, e1002246. 10.1371/journal.pcbi.1002246.

10. Del Alamo, D., Meiler, J., and McHaourab, H.S. (2022). Principles of Alternating Access in LeuT-fold Transporters: Commonalities and Divergences. J Mol Biol 434, 167746. 10.1016/j.jmb.2022.167746.

11. Perez, C., Koshy, C., Yildiz, O., and Ziegler, C. (2012). Alternating-access mechanism in conformationally asymmetric trimers of the betaine transporter BetP. Nature 490, 126–130. 10.1038/nature11403.

12. Holm, L. (2020). DALI and the persistence of protein shape. Protein Sci 29, 128–140. 10.1002/pro.3749.

13. 13. Zhao, Y., Shen, J., Wang, Q., Ruiz Munevar, M.J., Vidossich, P., De Vivo, M., Zhou, M., and Cao, E. (2022). Structure of the human cation-chloride cotransport KCC1 in an outward-open state. Proc Natl Acad Sci U S A 119, e2109083119. 10.1073/pnas.2109083119.

14. Ray, S., Berry, S.P., Wilson, E.A., Zhang, C.H., Shekhar, M., Singharoy, A., and Gaudet, R. (2023). High-resolution structures with bound Mn^2+^ and Cd^2+^ map the metal import pathway in an Nramp transporter. eLife 12, e84006. 10.7554/eLife.84006.

15. Nishikawa, K., Ooi, T., Isogai, Y., and Saitô, N. (1972). Tertiary Structure of Proteins. I. Representation and Computation of the Conformations. Journal of the Physical Society of Japan 32, 1331–1337. 10.1143/jpsj.32.1331.

16. Richards, F.M., and Kundrot, C.E. (1988). Identification of structural motifs from protein coordinate data: secondary structure and first-level supersecondary structure. Proteins 3, 71–84. 10.1002/prot.340030202.

17. Bozzi, A.T., Zimanyi, C.M., Nicoludis, J.M., Lee, B.K., Zhang, C.H., and Gaudet, R. (2019). Structures in multiple conformations reveal distinct transition metal and proton pathways in an Nramp transporter. eLife 8, e41124. 10.7554/eLife.41124.

18. Lomize, M.A., Pogozheva, I.D., Joo, H., Mosberg, H.I., and Lomize, A.L. (2012). OPM database and PPM web server: resources for positioning of proteins in membranes. Nucleic Acids Res 40, D370–376. 10.1093/nar/gkr703.

19. Gartner, R.M., Perez, C., Koshy, C., and Ziegler, C. (2011). Role of bundle helices in a regulatory crosstalk in the trimeric betaine transporter BetP. J Mol Biol 414, 327–336. 10.1016/j.jmb.2011.10.013.

20. Perez, C., Khafizov, K., Forrest, L.R., Kramer, R., and Ziegler, C. (2011). The role of trimerization in the osmoregulated betaine transporter BetP. EMBO Rep 12, 804–810. 10.1038/embor.2011.102.

21. Coleman, J.A., Yang, D., Zhao, Z., Wen, P.C., Yoshioka, C., Tajkhorshid, E., and Gouaux, E. (2019). Serotonin transporter-ibogaine complexes illuminate mechanisms of inhibition and transport. Nature 569, 141–145. 10.1038/s41586-019-1135-1.

22. Kazmier, K., Sharma, S., Quick, M., Islam, S.M., Roux, B., Weinstein, H., Javitch, J.A., and McHaourab, H.S. (2014). Conformational dynamics of ligand-dependent alternating access in LeuT. Nat Struct Mol Biol 21, 472–479. 10.1038/nsmb.2816.

23. Gao, X., Zhou, L., Jiao, X., Lu, F., Yan, C., Zeng, X., Wang, J., and Shi, Y. (2010). Mechanism of substrate recognition and transport by an amino acid antiporter. Nature 463, 828–832. 10.1038/nature08741.

24. Paz, A., Claxton, D.P., Kumar, J.P., Kazmier, K., Bisignano, P., Sharma, S., Nolte, S.A., Liwag, T.M., Nayak, V., Wright, E.M., et al. (2018). Conformational transitions of the sodium-dependent sugar transporter, vSGLT. Proc Natl Acad Sci U S A 115, E2742–E2751. 10.1073/pnas.1718451115.

25. Bozzi, A.T., Bane, L.B., Weihofen, W.A., McCabe, A.L., Singharoy, A., Chipot, C.J., Schulten, K., and Gaudet, R. (2016). Conserved methionine dictates substrate preference in Nramp-family divalent metal transporters. Proceedings of the National Academy of Sciences 113, 10310. 10.1073/pnas.1607734113.

26. Cui, W., Niu, Y., Sun, Z., Liu, R., and Chen, L. (2023). Structures of human SGLT in the occluded state reveal conformational changes during sugar transport. Nat Commun 14, 2920. 10.1038/s41467-023-38720-1.

27. Jumper, J., Evans, R., Pritzel, A., Green, T., Figurnov, M., Ronneberger, O., Tunyasuvunakool, K., Bates, R., Zidek, A., Potapenko, A., et al. (2021). Highly accurate protein structure prediction with AlphaFold. Nature 596, 583–589. 10.1038/s41586-021-03819-2.

28. del Alamo, D., Sala, D., McHaourab, H.S., and Meiler, J. (2022). Sampling alternative conformational states of transporters and receptors with AlphaFold2. eLife 11, e75751. 10.7554/eLife.75751.

29. Wayment-Steele, H.K., Ojoawo, A., Otten, R., Apitz, J.M., Pitsawong, W., Hömberger, M., Ovchinnikov, S., Colwell, L., and Kern, D. (2023). Predicting multiple conformations via sequence clustering and AlphaFold2. Nature. 10.1038/s41586-023-06832-9.

30. Tunyasuvunakool, K., Adler, J., Wu, Z., Green, T., Zielinski, M., Zidek, A., Bridgland, A., Cowie, A., Meyer, C., Laydon, A., et al. (2021). Highly accurate protein structure prediction for the human proteome. Nature 596, 590–596. 10.1038/s41586-021-03828-1.

31. Perland, E., and Fredriksson, R. (2017). Classification Systems of Secondary Active Transporters. Trends Pharmacol Sci 38, 305–315. 10.1016/j.tips.2016.11.008.

32. Lu, F., Li, S., Jiang, Y., Jiang, J., Fan, H., Lu, G., Deng, D., Dang, S., Zhang, X., Wang, J., and Yan, N. (2011). Structure and mechanism of the uracil transporter UraA. Nature 472, 243–246. 10.1038/nature09885.

33. Pei, J., Kim, B.-H., and Grishin, N.V. (2008). PROMALS3D: a tool for multiple sequence and structure alignment. Nucleic Acids Research 36, 2295–2300.

34. Virtanen, P., Gommers, R., Oliphant, T.E., Haberland, M., Reddy, T., Cournapeau, D., Burovski, E., Peterson, P., Weckesser, W., and Bright, J. (2020). SciPy 1.0: fundamental algorithms for scientific computing in Python. Nature methods 17, 261–272.

35. Waskom, M. (2021). seaborn: statistical data visualization. Journal of Open Source Software 6. 10.21105/joss.03021.

36. Pedregosa, F., Varoquaux, G., Gramfort, A., Michel, V., Thirion, B., Grisel, O., Blondel, M., Prettenhofer, P., Weiss, R., and Dubourg, V. (2011). Scikit-learn: Machine learning in Python. the Journal of machine Learning research 12, 2825–2830.

