## Supplementary Information for "They all rock: A systematic comparison of conformational movements in LeuT-fold transporters"

**Supplementary Note: Literature review**

Existing analyses of LeuT-fold proteins have identified four main conformational differences that occur during the transport cycle (Table S1 and references within): TM10a moves toward the hash and bundle to block the outer vestibule; the bundle rotates relative to the hash to close the outer vestibule and open the inner vestibule; TM5 moves away from the inner vestibule; and TM1a swings sharply up and away from the inner vestibule, possibly allowing substrate release. Each movement has been observed in at least three different LeuT-fold families to date. However, these four conformational movements are not uniform, and they can be more pronounced in certain proteins than others and can occur in different order, relative both to each other and to substrate binding and release.

**The $O_{op} \rightarrow O_{oc}$ transition involves movement of TM6a and TM10a**

The $O_{op} \rightarrow O_{oc}$ transition occurs after the binding of substrate (and any co-substrate) in importers[1]. During this transition, TM6a (in AdiC and SERT) or TM10a (in AdiC, SERT, Mhp1, and in MntH for the $O_{op} \rightarrow I_{oc}$ comparison) may move into the outer vestibule and rotate toward the membrane plane (Table S1)[2-4]. For Mhp1, TM10a moves toward the outer bundle in both the $O_{op} \rightarrow O_{oc}$ and $O_{oc} \rightarrow I_{op}$ comparisons, suggesting that its movement can span multiple transitions[2-4]. In all $O_{oc}$ structures, the binding site is blocked either by TM6a's proximity to the hash or TM10's interaction with TM1b, explaining the necessity of movement by TM6a or TM10a (Table S2).

**The outward-to-inward transition involves bundle-hash movement and movement of arms**

The outward-to-inward transition involves a continuation of the movements from the $O_{op} \rightarrow O_{oc}$ transition that close the outer vestibule as well as movements to open the inner vestibule. In all

LeuT-fold transporters with structures in multiple conformations, except BetP, previous analyses have noted that the bundle tilts relative to the hash, closing the outer vestibule and opening the inner vestibule (Table S1). During this movement, the angle between the bundle helices and TM4 of the hash decreases[5]. In LeuT, SERT, Mhp1, and LAT1, the entire bundle moves toward the hash in the outer vestibule and away from the hash in the inner vestibule[3,6-8]. In contrast, TM6a is the only bundle helix that has been reported to move independently during the outward-to-inward transition, in MntH[9]. In BetP, the outward-to-inward transition has been described as the outer portion of TM3 moving closer to and the inner portion of TM9 moving away from the bundle instead of the bundle moving relative to the hash[10]. Similarly, in KCC1 rocking of TM3 and TM8, as well as movement of TM4 and TM9, allow for alternating access[11]. The bundle can move either like a rigid body, like in Mhp1 where the $I_{op}$ structure superimposes almost exactly with a model based on a screw transformation of the bundle from the $O_{op}$ structure[12] or with independent helix movements, such as in MntH[9].

The outward-to-inward transition also involves movement of TM5 away from the inner hash to open the inner vestibule, as seen in SERT, LeuT, Mhp1, and MntH (Table S1). TM5 movement is also seen in the $I_{oc}{\rightarrow}I_{op}$ comparisons for SGLT and LeuT, and in $O_{op}{\rightarrow}I_{oc}$ for MntH (Table S1). The TM5 movement is greater in Mhp1 than in SGLT and LeuT (Table S1). The movement of TM5 away from the inner vestibule in SERT was proposed to allow $Na^+$ release[3].

**The $I_{oc}{\rightarrow}I_{op}$ transition involves swinging of TM1a**

Finally, the $I_{oc}{\rightarrow}I_{op}$ transition involves an independent swinging movement of TM1a up and away from the bundle in LeuT, SGLT, SERT, and MntH (Table S1)[3,5,6,13]. Deletion of TM1a interfered with transport in the *Deinococcus radiodurans* homolog of MntH[13] but did not completely abrogate transport in the *Staphylococcus capitis* homolog[14]. In site-directed spin labelling experiments with LeuT, TM1a did not show as large a magnitude of movement as expected from the structural comparison of the $O_{oc} \rightarrow I_{op}$ structures, leading to the suggestion that the structural data may overestimate this movement[6].

**Table S1. Conformational movements of described in literature.** Unless otherwise noted, the outward to inward transition is described.

| Outer bundle helices | | | | | |
|---|---|---|---|---|---|
| **Protein** | **TM1b** | **TM2a** | **TM6a** | **TM7b** | **Refs** |
| AdiC | | moves toward the hash during $O_{op} \rightarrow O_{oc}$ transition | moves (more than other helices) toward the hash to occlude the binding site from the outer vestibule | | 2 |
| BetP | bundle undergoes little movement throughout transport cycle | | | | 10 |
| KCC1 | bundle undergoes little movement throughout transport cycle | | | | 11 |
| LAT1 | bundle rotates 27º relative to hash | | | | 8 |
| LeuT | moves toward TM3 and TM5 of the scaffold | | moves toward TM3 and TM5 of the scaffold | moves toward TM3 and TM5 of the scaffold | 6 |
| Mhp1 | bundle and hash undergo little movement during outward occlusion, but hash rotates 30º with respect to the bundle as a rigid body during outward closing | | | | 7 |
| MntH | compared to other transporters, less movement of outer bundle helices, and bundle and hash do not move as strict rigid bodies | | only mobile helix among bundle helices | | 9 |
| SERT | rotates 22° and translates 5.1 Å towards the hash | rotates 7.3° and translates 2.8 Å towards the hash | rotates 3° and translates 1.9 Å towards the hash during the $O_{op} \rightarrow O_{oc}$ transition; rotates 5° and translates 3.4 Å towards the hash during the $O_{oc} \rightarrow I_{op}$ transition | rotates 4.8° and translates 1.0 Å towards the hash | 3 |

| Inner bundle helices | | | | | |
|---|---|---|---|---|---|
| **Protein** | **TM1a** | **TM2b** | **TM6b** | **TM7a** | |
| BetP | bundle undergoes little movement throughout transport cycle | | | | 10 |
| | inner vestibule in $I_{op}$ is narrower than in other transporters; negatively charged membrane lipid restricts the movement of the bundle | | | | 15 |
| KCC1 | bundle undergoes little movement throughout transport cycle | | | | 11 |
| LAT1 | | hydrogen bond with TM10b is broken | | | 8 |
| LeuT | rotates 11º in plane of membrane from $O_{oc} \rightarrow I_{oc}$; 136º kink in $O_{oc}$ and 125º kink in $I_{op}$; N-terminus of TM1a unwind from $I_{oc} \rightarrow I_{op}$ | | moves upon substrate binding to open inner vestibule; Y268 necessary for inward-closing movement of TM5 | moves away from TM4 and intracellular loop between TM2 and TM3 | 6,9,16 |
| Mhp1 | hash rotates 30º with respect to the bundle as a rigid body during outward closing | | | | 7 |
| MntH | 103º kink in $I_{op}$ | mostly stationary during transport | | | 9,13 |
| SERT | rotates 40° in plane of membrane, which disrupts interactions between the N terminus and TM6b that occur in outward conformations | | | | 5 |
| SGLT | kinks outward during transition to $I_{op}$ | | | | 5 |

| TM5 and TM10 | | | |
|---|---|---|---|
| **Protein** | **TM5** | **TM10** | **Refs** |

| | | | |
|---|---|---|---|
| AdiC | | moves inward to obstruct the outer vestibule | 2 |
| LAT1 | | hydrogen bond with TM2 breaks; inhibitor blocks the rotation of TM10 in $O_{op}$ conformation | 8 |
| LeuT | angle between TM5 and membrane plane decreases by 30.0º | | 5 |
| | responsible for inward closing movement | | 6 |
| | loop between TM4 and TM5 extends, so TM5 tilts away from TM1 from $O_{oc} \rightarrow I_{oc}$, which solvates $Na^+$ binding site | | 16 |
| Mhp1 | angle between TM5 and membrane plane decreases by 24.1º from $O_{oc} \rightarrow I_{op}$; rotates out and away from the cytosolic pathway | angle between TM10 and membrane plane increases in total by 36º (roughly equally during the $O_{op} \rightarrow O_{oc}$ and outward to inward steps); moves inward to obstruct the outer vestibule | 5 |
| | less flexible when outward conformation stabilized | flexibility increases when outward conformation stabilized | 4 |
| MntH | shows large movement | outer half shows large movement | 9 |
| SERT | rotates 3.2° and translates 1.3 Å toward the bundle during outward occlusion; rotates 7º from outward to inward; TM5a and TM5b translate 1.8 Å and 1.0 Å away from the binding site, respectively | rotates 2° and translates 0.9 Å towards the bundle during outward occlusion | 3 |
| SGLT | movement away from hash facilitates inward closing, but less in SGLT and LeuT than in Mhp1 | predicted to move 16 Å to obstruct the outer vestibule | 17 |

62

**Table S2. Literature-based identification of the helices or sidechains that occlude the binding site in each occluded structure.** Occlusion can be mediated by entire helices or by a single sidechain.

| Conformation | Protein | Structure | What occludes binding site? | Refs |
|---|---|---|---|---|
| $O_{oc}$ | AdiC | 3L1L | TM6a moves, displacing the W202 sidechain 10 Å from $O_{op} \rightarrow O_{oc}$, which blocks outer vestibule[1] | 2 |
| $O_{oc}$ | BetP | 4AIN | W377 (TM6a) rotates by 90º to block outer vestibule[2] | 10 |
| $O_{oc}$ | Mhp1 | 4D1B | TM10b folds over the binding site to occlude substrate[3] | 18 |
| $O_{oc}$ | SERT | 6DZV | Movement of TM6a toward scaffold occludes binding site[4] | 3 |
| $O_{oc}$ | LeuT | 2A65 | Substrate is directly occluded from outer vestibule by sidechains of Y108 (TM3) and F253 (TM6a), and above those sidechains is an ion pair of R30 (TM1b) and D404 (TM10a)[5] | 19 |
| $I_{oc}$ | LeuT | 6XWM | Y268 (TM6b), Q361, and D369 (both TM8) occlude the inner vestibule[6] | 16 |
| $I_{oc}$ | SGLT | 3DH4 | M73, Y87 (both TM1a), and F424 (TM8) occlude the substrate from the inner vestibule[7] | 20 |
| $I_{oc}$ | *Dr*MntH | 6C3I | TM1a moves away from TM8 to solvate the binding site from the interior aqueous vestibule[8] | 21 |

**Table S3. List of available comparisons**

| | Conformations | | | | Comparisons | | | | | |
|---|---|---|---|---|---|---|---|---|---|---|
| **Protein** | $\mathbf{O_{op}}$ | $\mathbf{O_{oc}}$ | $\mathbf{I_{oc}}$ | $\mathbf{I_{op}}$ | $\mathbf{O_{op} \rightarrow I_{op}}$ | $\mathbf{O_{op} \rightarrow O_{oc}}$ | $\mathbf{O_{op} \rightarrow I_{oc}}$ | $\mathbf{O_{oc} \rightarrow I_{oc}}$ | $\mathbf{O_{oc} \rightarrow I_{op}}$ | $\mathbf{I_{oc} \rightarrow I_{op}}$ |
| *Ec*AdiC | X | X | | | | X | | | | |
| *Cg*BetP | X | X | | X | X | X | | | X | |
| *Dr*MntH | X | | X | X | X | | X | | | X |
| *Hs*SERT | X | X | | X | X | X | | | X | |
| *Aa*LeuT | X | X | X | X | X | X | X | X | X | X |
| *Ml*Mhp1 | X | X | | X | X | X | | | X | |
| *Vp*SGLT | | | X | X | | | | | | X |
| *Hs*KCC1 | X | | | X | X | | | | | |
| *Hs*LAT1 | X | | | X | X | | | | | |
| **Total** | **8** | **5** | **3** | **8** | **7** | **5** | **2** | **1** | **4** | **3** |

**Table S4. Transmembrane helix ranges determined for each protein**

| TM | LeuT | SERT | BetP | Mhp1 | MntH | KCC1 | LAT1 | AdiC | SGLT |
|---|---|---|---|---|---|---|---|---|---|
| 1a | 13-21 | 85-96 | 139-147 | 30-41 | 46-54 | 121-133 | 53-64 | 13-24 | 54-65 |
| 1b | 27-38 | 103-111 | 152-168 | 43-55 | 57-70 | 137-147 | 67-76 | 27-36 | 67-80 |
| 2 | 43-69 | 117-141 | 177-210 | 59-86 | 74-100 | 148-174 | 86-108 | 43-65 | 82-109 |
| 3 | 88-121 | 159-185 | 235-266 | 103-134 | 114-146 | 196-225 | 127-153 | 84-109 | 125-158 |
| 4 | 167-184 | 255-269 | 280-292 | 142-159 | 150-165 | 250-268 | 171-188 | 125-142 | 161-176 |
| 5a | 191-198 | 279-288 | 301-306 | 161-174 | 175-182 | 272-282 | 192-202 | 146-156 | 187-198 |
| 5b | 199-209 | 289-299 | 312-324 | 175-186 | 183-194 | 283-293 | 203-214 | 157-168 | 199-210 |
| 6a | 241-254 | 325-337 | 366-373 | 208-220 | 216-227 | 419-428 | 242-251 | 192-201 | 249-264 |
| 6b | 260-267 | 342-350 | 379-387 | 222-232 | 231-236 | 434-440 | 258-264 | 208-214 | 269-276 |
| 7 | 276-306 | 360-385 | 394-426 | 246-278 | 257-286 | 448-471 | 271-294 | 221-244 | 280-312 |
| 8 | 338-370 | 422-450 | 452-479 | 296-329 | 311-340 | 504-532 | 325-352 | 275-303 | 349-380 |
| 9 | 375-396 | 464-479 | 492-506 | 337-349 | 352-365 | 555-569 | 374-388 | 324-339 | 394-414 |
| 10a | 398-407 | 487-495 | 515-524 | 359-369 | 370-383 | 576-583 | 395-402 | 352-359 | 422-432 |
| 10b | 409-423 | 500-513 | 527-538 | 370-382 | 384-396 | 584-597 | 403-416 | 360-373 | 434-447 |

**Table S5. Structures used as inputs for DALI searches**

| Protein | Structure (PDB ID including chain) | Hits |
|---|---|---|
| AdiC | 5J4NA | 246 |
| ApcT | 3GIAA | 298 |
| BetP | 4DOJA | 296 |
| CaiT | 2WSWA | 297 |
| DAT | 4XPFA | 297 |
| GadC | 4DJKA | 298 |
| LeuT | 2Q6HA | 298 |
| Mhp1 | 4D1DA | 272 |
| MhsT | 4US3A | 271 |
| MntH | 6C3IA | 296 |
| SERT | 6AWNA | 238 |
| SGLT | 3DH4A | 296 |

**Table S6. Statistics of RMSDD value distributions for intrahelical and interhelical comparisons.**

| | | | K-S test statistics | | | | | |
|---|---|---|---|---|---|---|---|---|
| | | | Intrahelical vs. intrahelical | | Interhelical vs. intrahelical | | Interhelical vs. interhelical | |
| Helix | Mean RMSDD (Å) | Standard deviation (Å) | *statistic* | *p-value* | *statistic* | *p-value* | *statistic* | *p-value* |
| 1a | 0.503 | 0.381 | 0.262 | 0.103 | 0.558 | 6.35E-07 | 0.245 | 1.88E-14 |
| 1b | 0.323 | 0.174 | 0.129 | 0.848 | 0.682 | 1.27E-10 | 0.08 | 0.063 |
| 2 | 0.351 | 0.195 | 0.171 | 0.539 | 0.667 | 4.23E-10 | 0.116 | 0.002 |
| 3 | 0.389 | 0.195 | 0.189 | 0.417 | 0.675 | 2.31E-10 | 0.096 | 0.014 |
| 4 | 0.427 | 0.503 | 0.136 | 0.802 | 0.61 | 2.46E-08 | 0.065 | 0.199 |
| 5a | 0.447 | 0.367 | 0.168 | 0.565 | 0.507 | 9.92E-06 | 0.184 | 2.51E-08 |
| 5b | 0.355 | 0.31 | 0.252 | 0.129 | 0.711 | 1.16E-11 | 0.102 | 0.007 |
| 6a | 0.468 | 0.399 | 0.136 | 0.802 | 0.539 | 1.79E-06 | 0.071 | 0.132 |
| 6b | 0.325 | 0.219 | 0.175 | 0.514 | 0.673 | 2.56E-10 | 0.114 | 0.002 |
| 7 | 0.458 | 0.242 | 0.378 | 0.004 | 0.575 | 2.33E-07 | 0.103 | 0.006 |
| 8 | 0.43 | 0.189 | 0.315 | 0.028 | 0.62 | 1.24E-08 | 0.1 | 0.009 |
| 9 | 0.333 | 0.259 | 0.199 | 0.352 | 0.68 | 1.51E-10 | 0.055 | 0.395 |
| 10a | 0.482 | 0.587 | 0.164 | 0.592 | 0.594 | 7.13E-08 | 0.214 | 4.31E-11 |
| 10b | 0.245 | 0.13 | 0.325 | 0.021 | 0.779 | 1.57E-14 | 0.158 | 2.93E-06 |

**Table S7. Angles between predicted membrane planes of transporter structures in $O_{op}$ and $I_{op}$ conformations when aligning by distinct structural element**

| Protein | Number of LeuT-fold protomers in native oligomer | Bundle | | Hash | | Core | |
|---|---|---|---|---|---|---|---|
| | | RMSD (Å) | θ (°) | RMSD (Å) | θ (°) | RMSD (Å) | θ (°) |
| LeuT | monomer[22] | 3.1 | 19 | 0.7 | 4 | 3.4 | 7 |
| SERT | monomer[23] | 2.7 | 16 | 1.0 | 4 | 2.8 | 9 |
| BetP | trimer[24] | 1.2 | 5 | 1.6 | 9 | 2.1 | 1 |
| Mhp1 | monomer[25] | 0.6 | 12 | 0.9 | 15 | 3.1 | 3 |
| MntH | monomer[14] | 2.2 | 8 | 0.9 | 5 | 3.0 | 3 |
| KCC1 | monomer[a] | 0.9 | 11 | 0.7 | 12 | 2.3 | 5 |
| LAT1 | monomer[26] | 2.1 | 17 | 1.2 | 11 | 3.3 | 7 |

[a]KCC1 can adopt a monomeric or dimeric state under native conditions[27]. Monomer was used because PPM3.0 cannot fit both protomers of the dimeric $O_{op}$ structure (PDB: 7TTI) into a planar membrane.

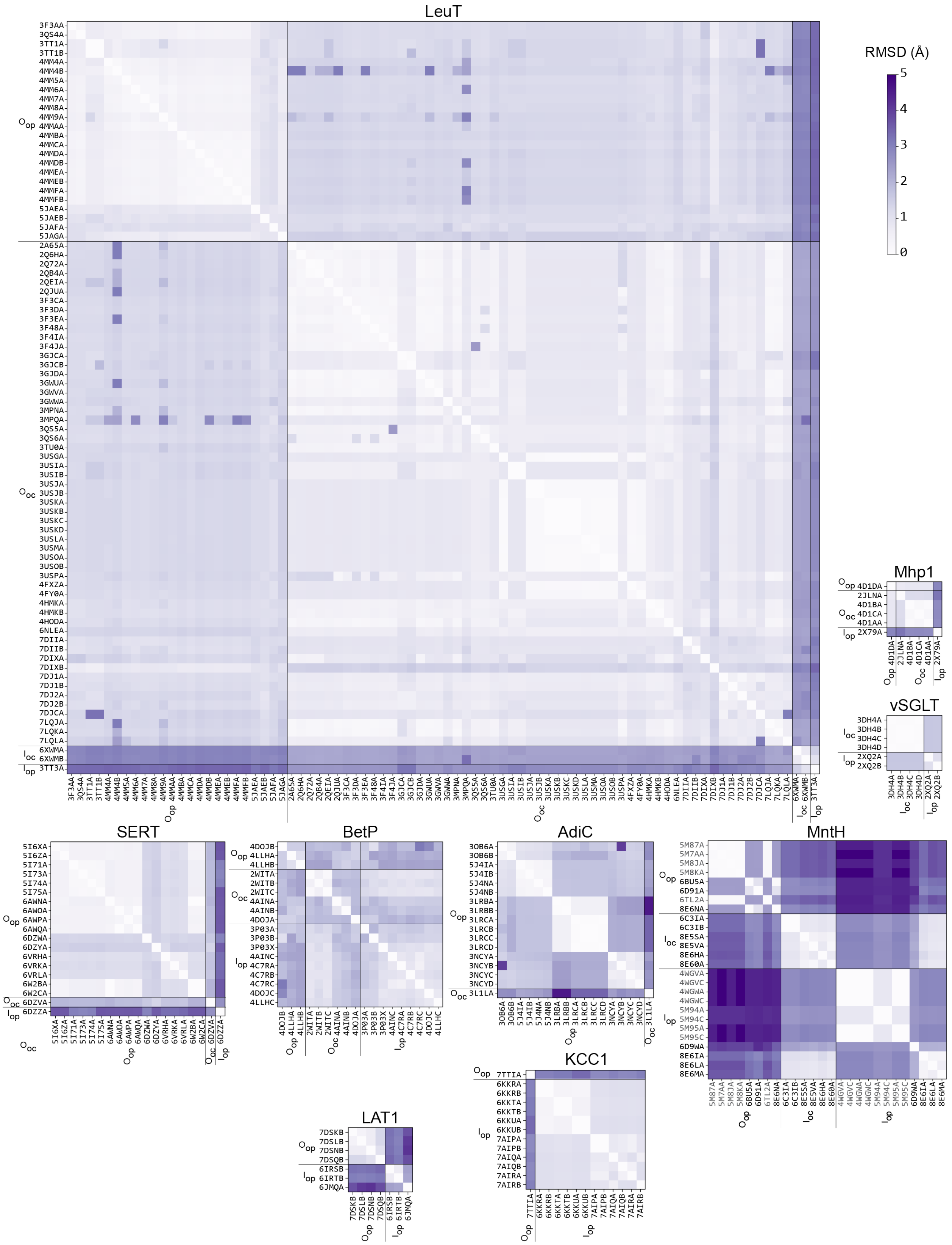


**Figure S1. RMSD matrices for LeuT-fold transporters with structures in more than one conformation.**

Conformations were assigned through a combination of clustering of similar structures and information gathering from the relevant publications. For KCC1, the RMSD matrices were constructed based on alignment of its transporter domain (omitting the C-terminal cytosolic domain), i.e., up to residue 655. In the case of MntH, PDB IDs in black correspond to *Deinococcus radiodurans* (Dra)MntH, and PDB IDs in grey to *Staphylococcus capitis* or *Eremococcus coleocola* MntH.

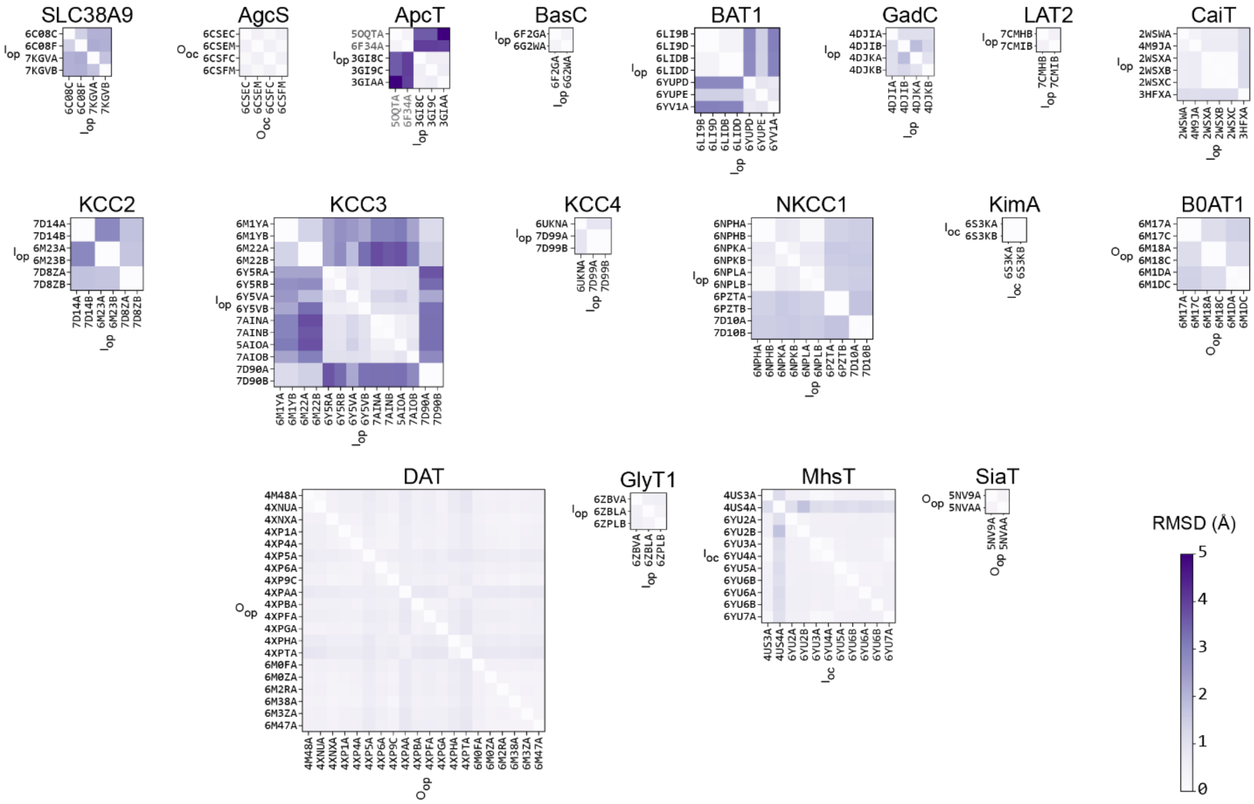


**Figure S2. RMSD matrices for LeuT-fold transporters with multiple structures assigned to the same conformation.**

Conformations were assigned from information gathering from the relevant publications and visual confirmation. For the CCC-family transporters, the RMSD matrices were constructed based on alignment of their transporter domains (omitting their C-terminal cytosolic domains), i.e., up to residue 660 (KCC2 and KCC4), or 675 (KCC3 and NKCC1). In the case of ApcT, PDB ID codes in grey correspond to *Geobacillus kaustophilus* ApcT, and those in black to *Methanocaldococcus jannaschii*.

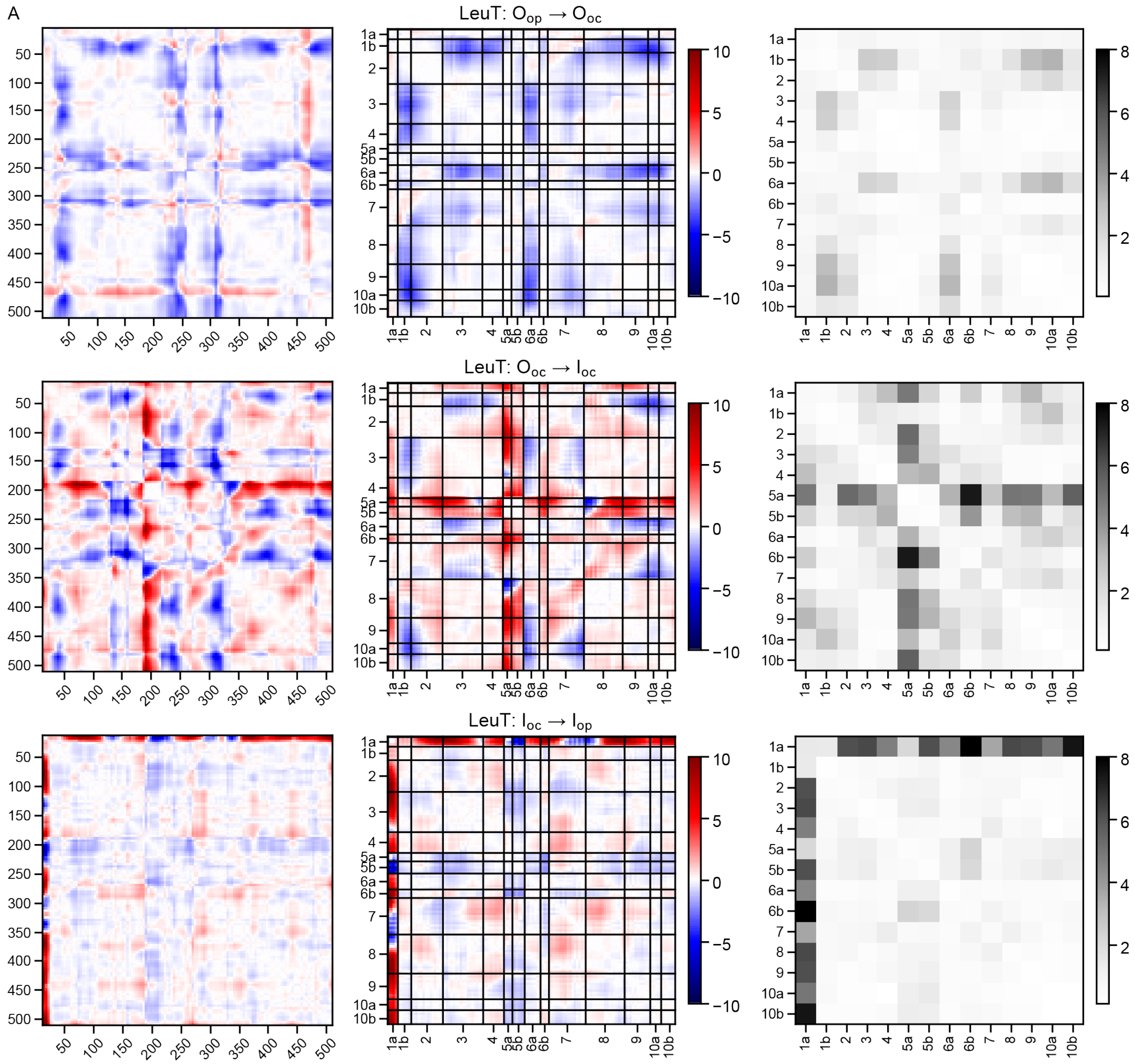


**Figure S3. All DDM matrices for the 22 pairwise structural comparisons (this page and the following 6 pages).**
(A) DDM matrices for LeuT one-step transitions. (B) DDM matrices for LeuT two-step and three-step transitions. (C) DDM matrices for SERT. (D) DDM matrices for BetP. (E) DDM matrices for Mhp1. (F) DDM matrices for MntH. (G) DDM matrices for KCC1. (H) DDM matrices for LAT1. (I) DDM matrices for AdiC. (J) DDM matrices for SGLT.

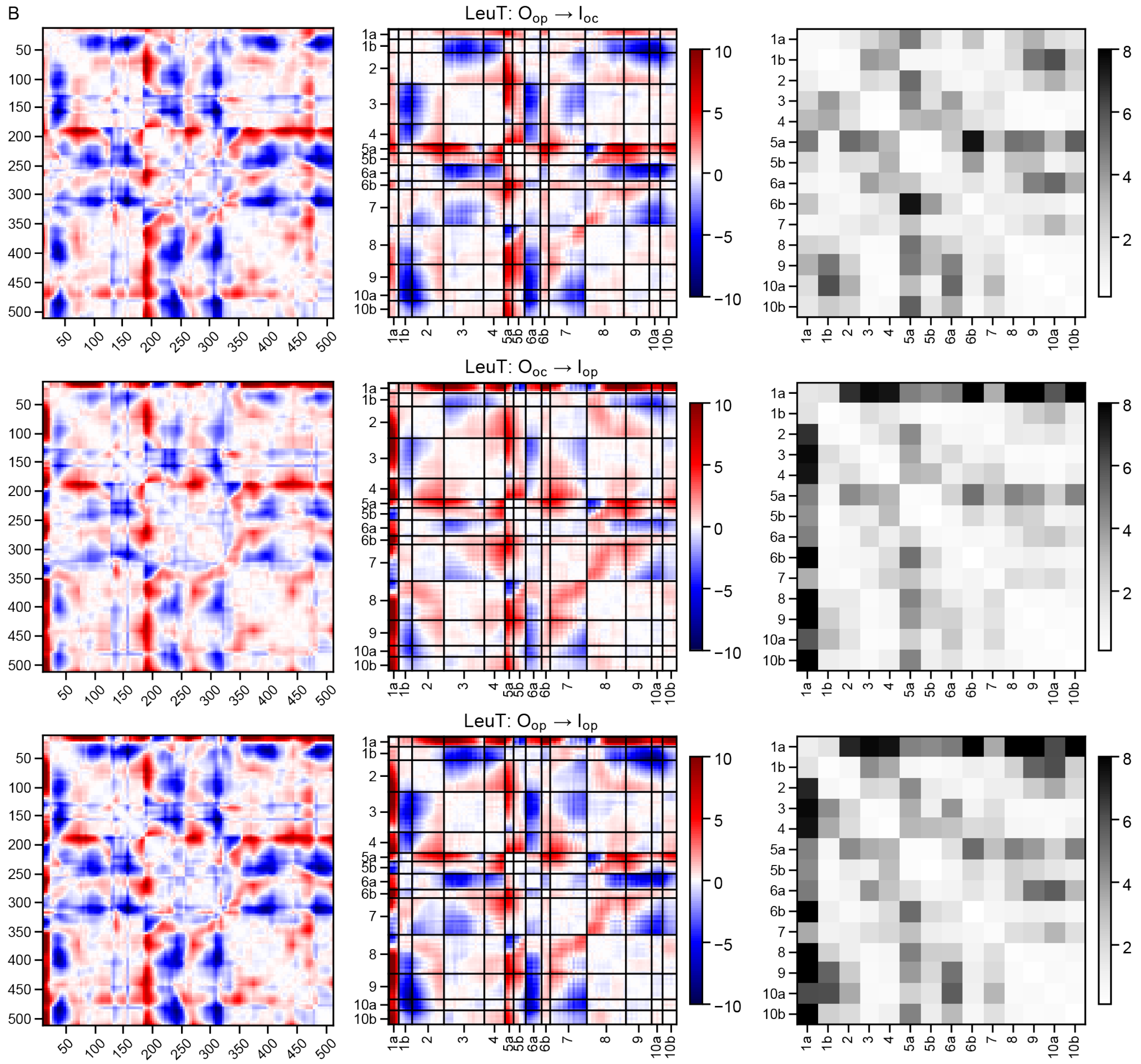
B
LeuT: O$_{op}$ → I$_{oc}$
LeuT: O$_{oc}$ → I$_{op}$
LeuT: O$_{op}$ → I$_{op}$

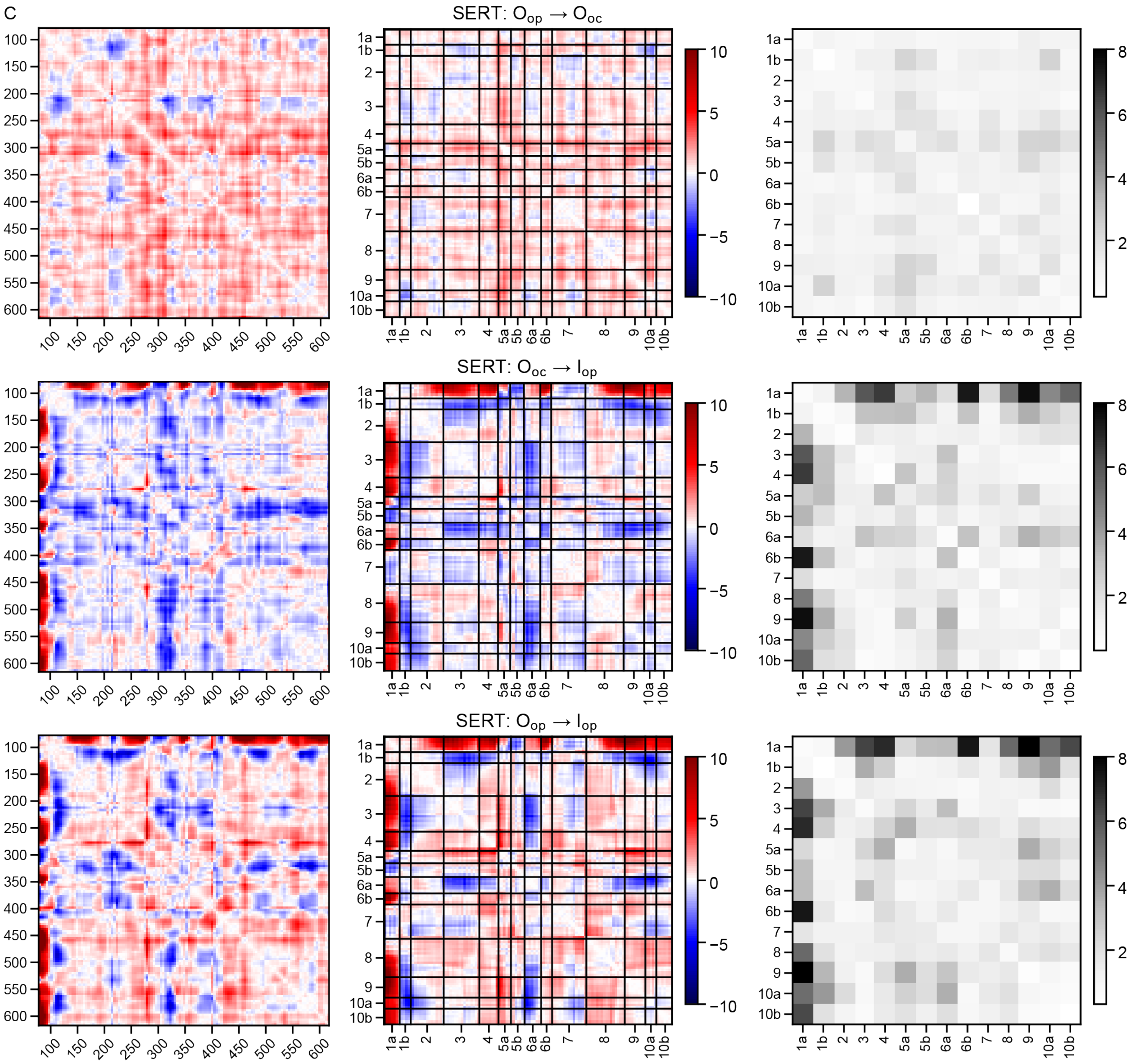
C
SERT: $O_{op} \rightarrow O_{oc}$
SERT: $O_{oc} \rightarrow I_{op}$
SERT: $O_{op} \rightarrow I_{op}$
1a
1b
2
3
4
5a
5b
6a
6b
7
8
9
10a
10b

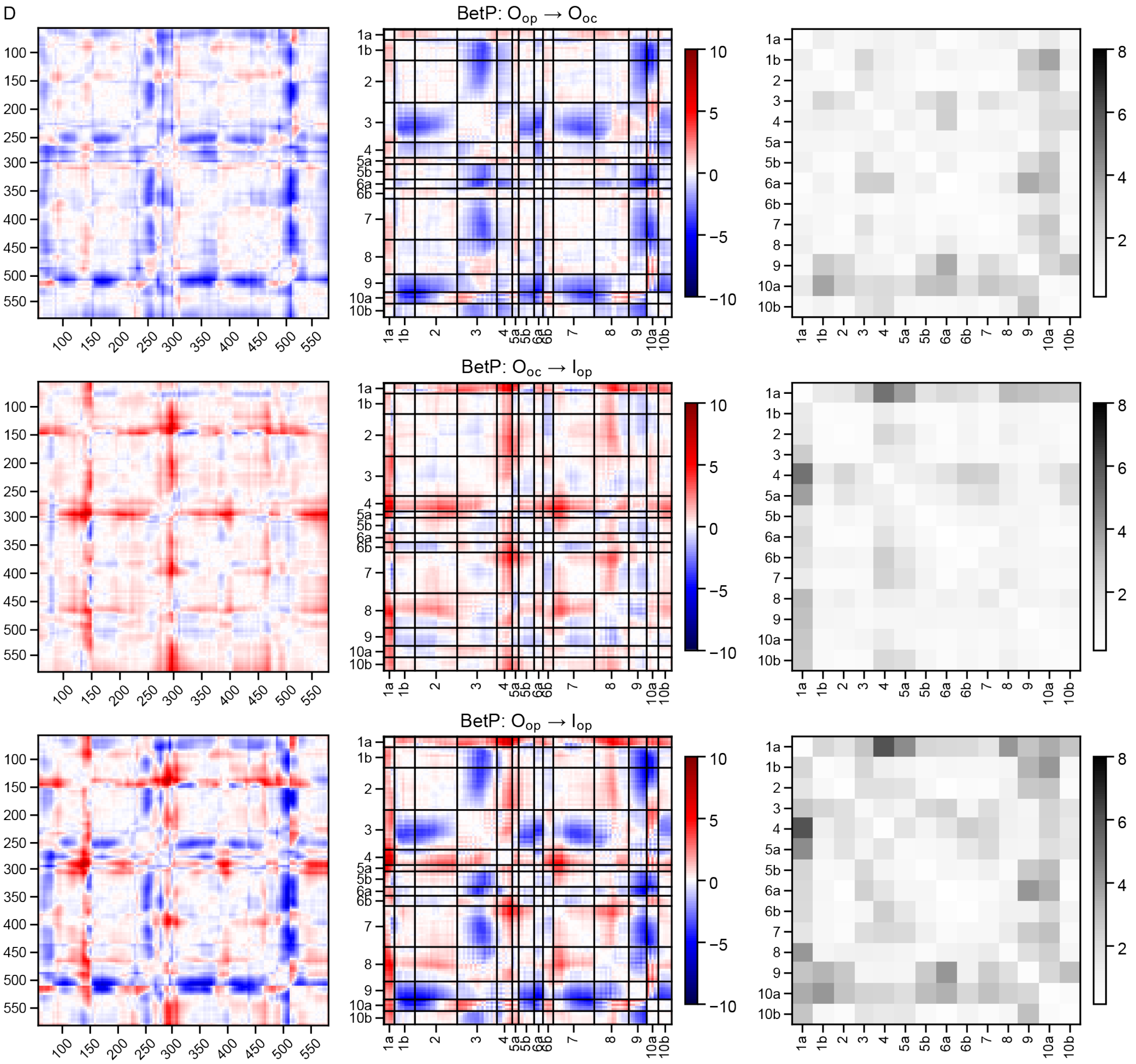
D
BetP: $O_{op} \rightarrow O_{oc}$
BetP: $O_{oc} \rightarrow I_{op}$
BetP: $O_{op} \rightarrow I_{op}$
100
150
200
250
300
350
400
450
500
550
1a
1b
2
3
4
5a
5b
6a
6b
7
8
9
10a
10b
10
5
0
−5
−10
8
6
4
2

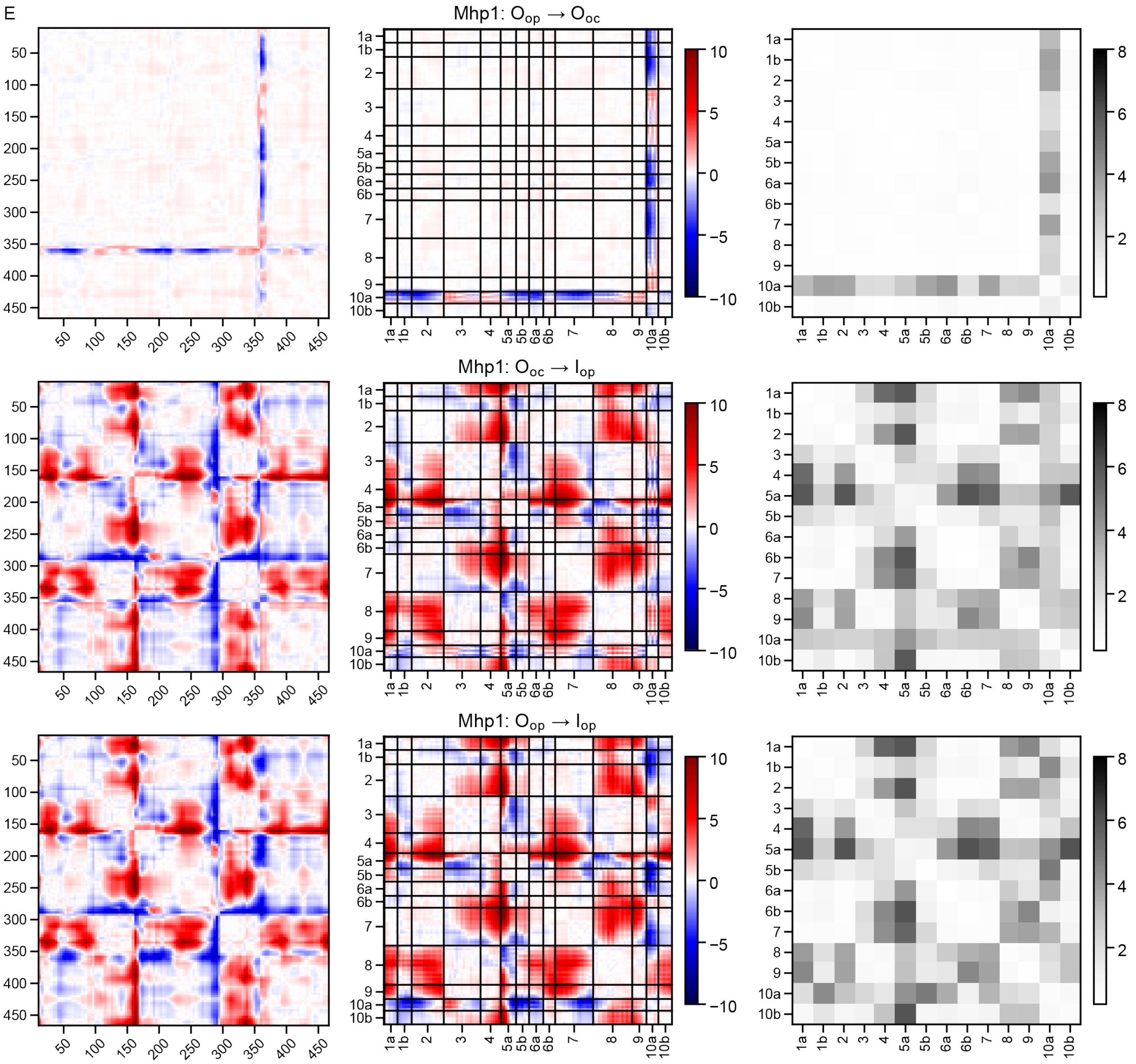
E
Mhp1: O$_{op}$ → O$_{oc}$
Mhp1: O$_{oc}$ → I$_{op}$
Mhp1: O$_{op}$ → I$_{op}$
1a 1b 2 3 4 5a 5b 6a 6b 7 8 9 10a 10b
50 100 150 200 250 300 350 400 450
10 5 0 −5 −10
8 6 4 2

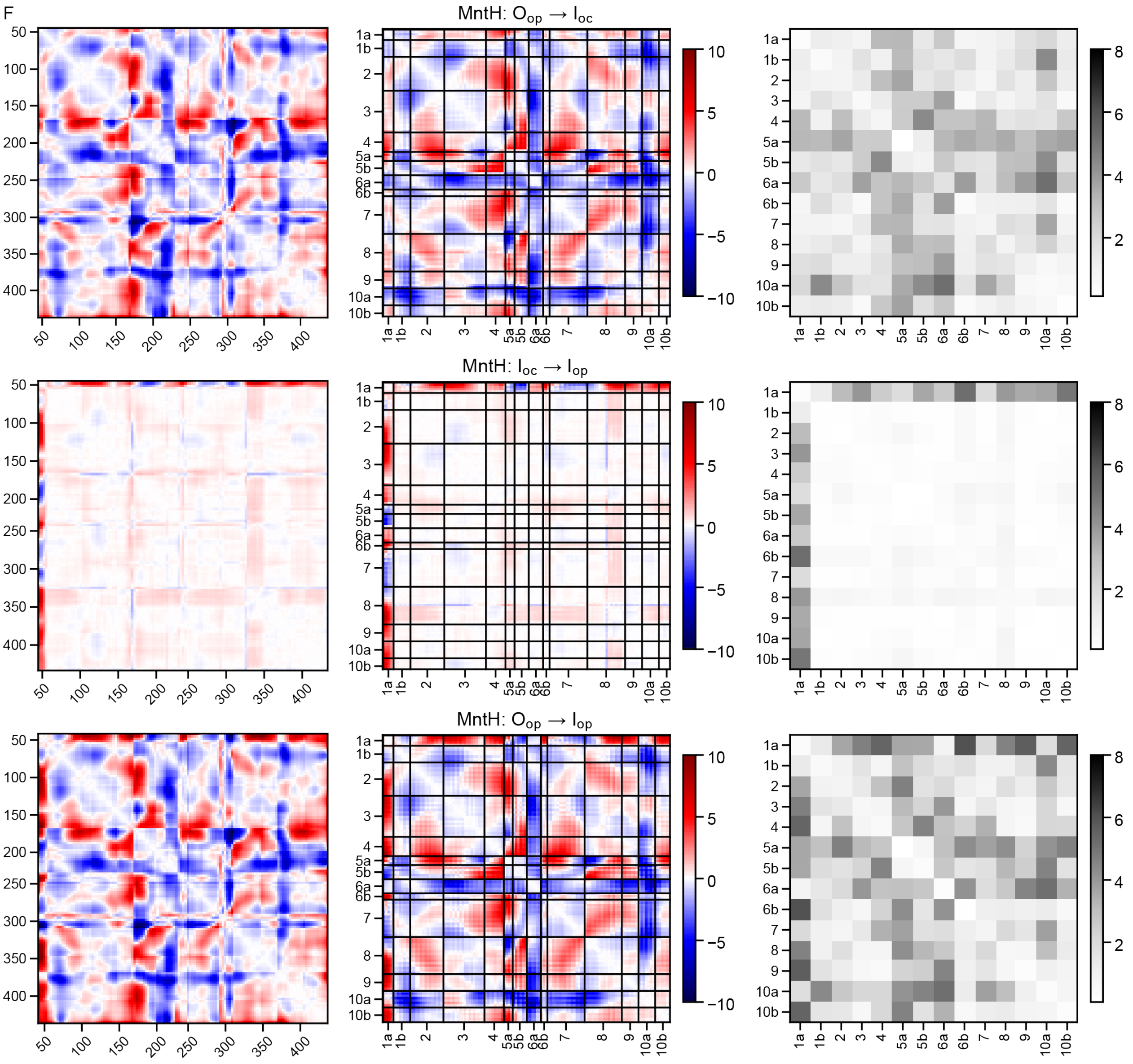
F
MntH: O$_{op}$ → I$_{oc}$
MntH: I$_{oc}$ → I$_{op}$
MntH: O$_{op}$ → I$_{op}$
1a
1b
2
3
4
5a
5b
6a
6b
7
8
9
10a
10b
10
5
0
−5
−10
8
6
4
2
50
100
150
200
250
300
350
400

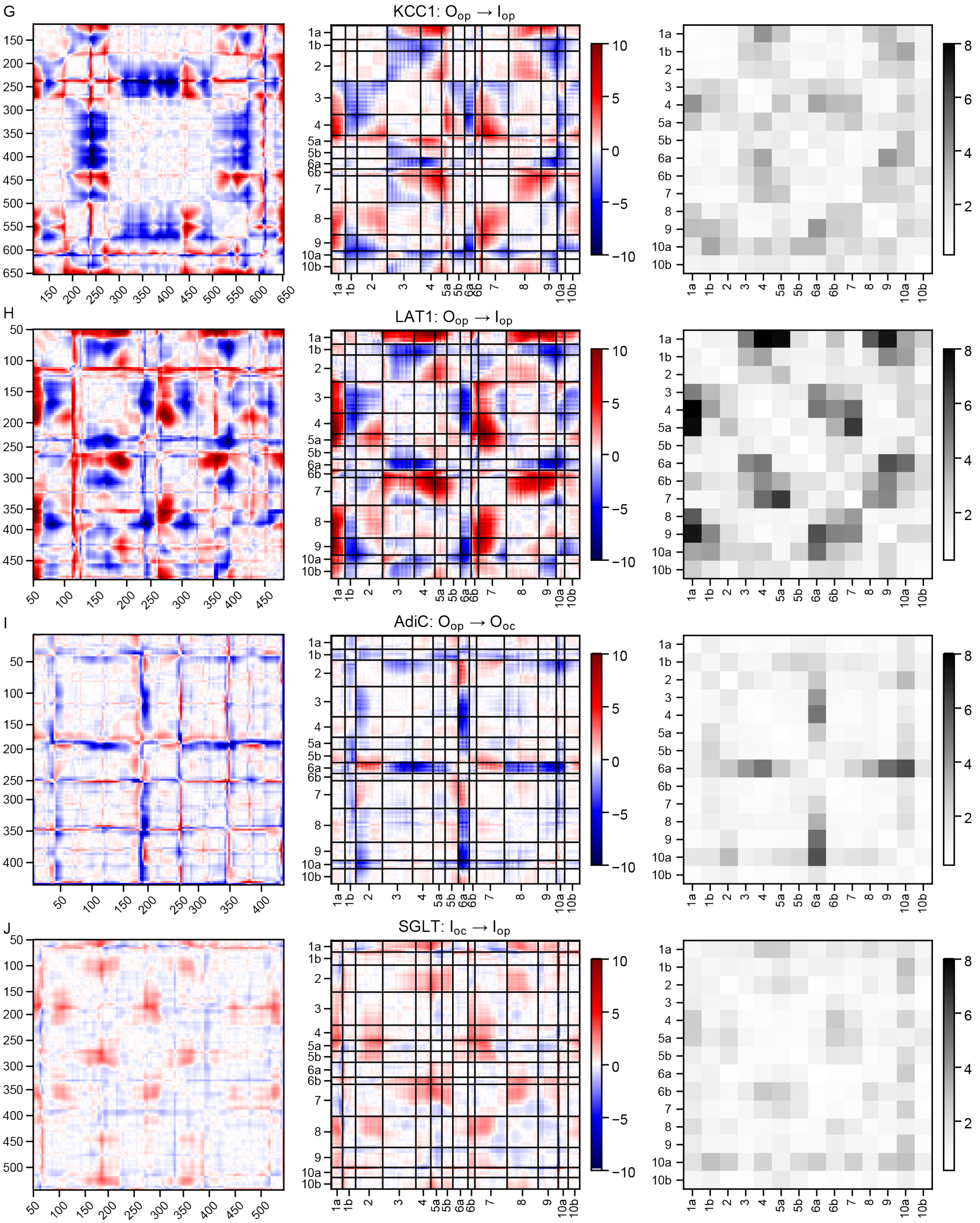
G
KCC1: $O_{op} \rightarrow I_{op}$
H
LAT1: $O_{op} \rightarrow I_{op}$
I
AdiC: $O_{op} \rightarrow O_{oc}$
J
SGLT: $I_{oc} \rightarrow I_{op}$

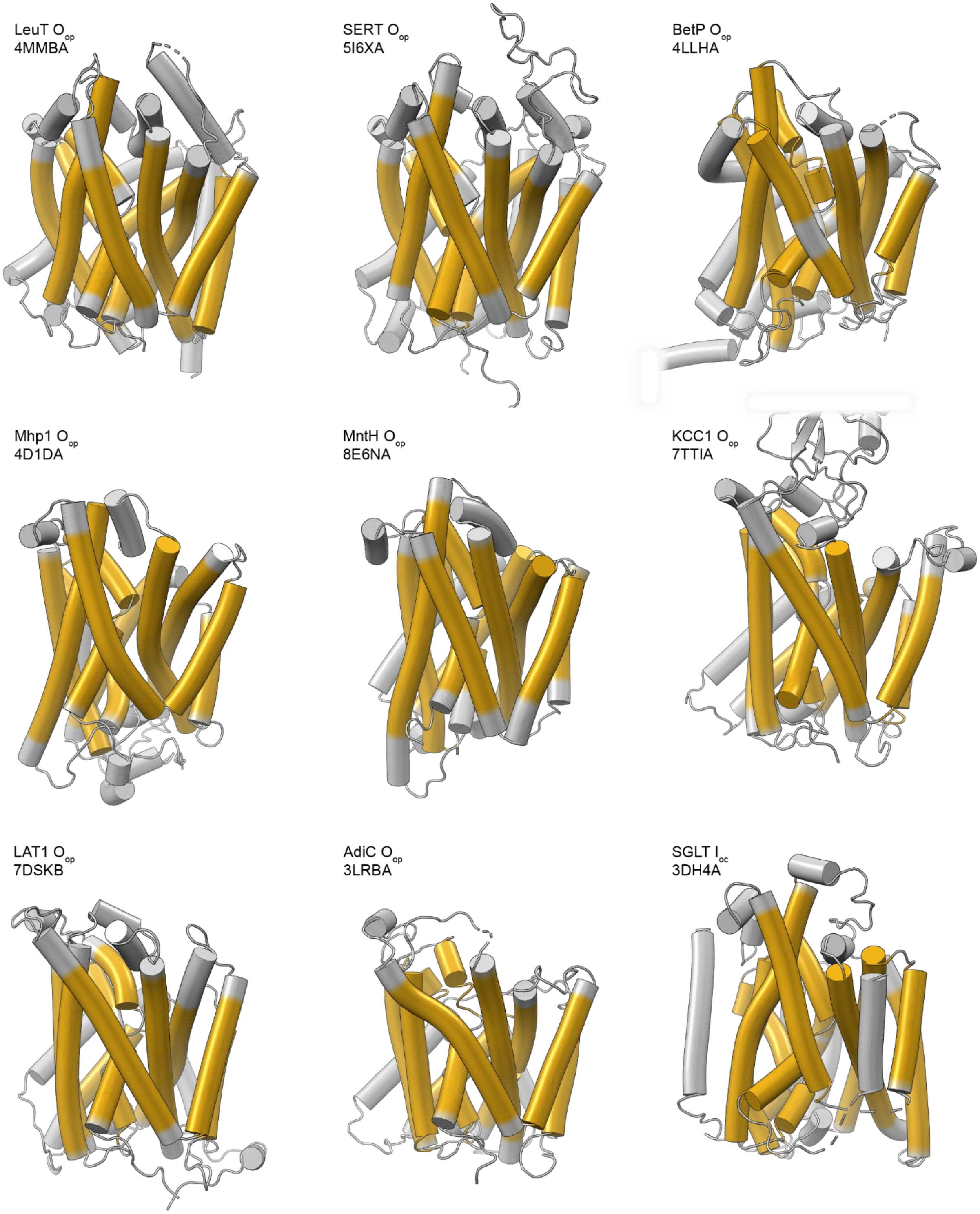


**Figure S4. Illustrations of the helical residue ranges used in our analyses (yellow) for the nine proteins analyzed.**

The $O_{op}$ structures are illustrated except for SGLT, for which there is no $O_{op}$ structure and the $I_{oc}$ structure is illustrated instead. The C-terminal helix of BetP and extracellular domain of KCC1 are truncated in this view.

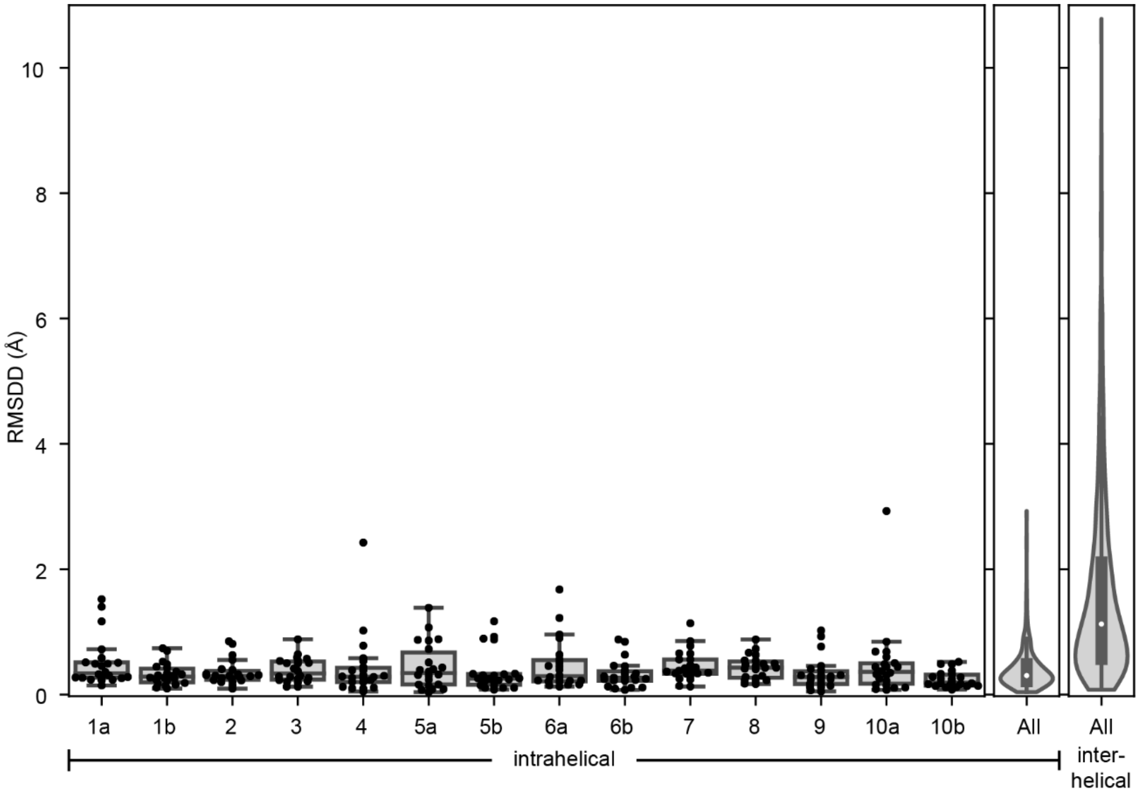


**Figure S5. Intrahelix root-mean-square distance difference (RMSDD) values validate the selection of transmembrane helix ranges.**

The distributions of RMSDDs for each of the 14 helices with itself (intrahelical distance differences) for all twenty-two comparisons are shown on the left, with violin plots on the right representing all intrahelical distance differences or interhelical RMSDDs. No individual helix has a distribution of intrahelical RMSDDs consistent with the interhelical RMSDDs, which would suggest that it should be subdivided. The two largest outliers are TM10a during the $I_{oc}$→$I_{op}$ transition of SGLT and TM4 during the $O_{op}$→$I_{op}$ transition of SERT.

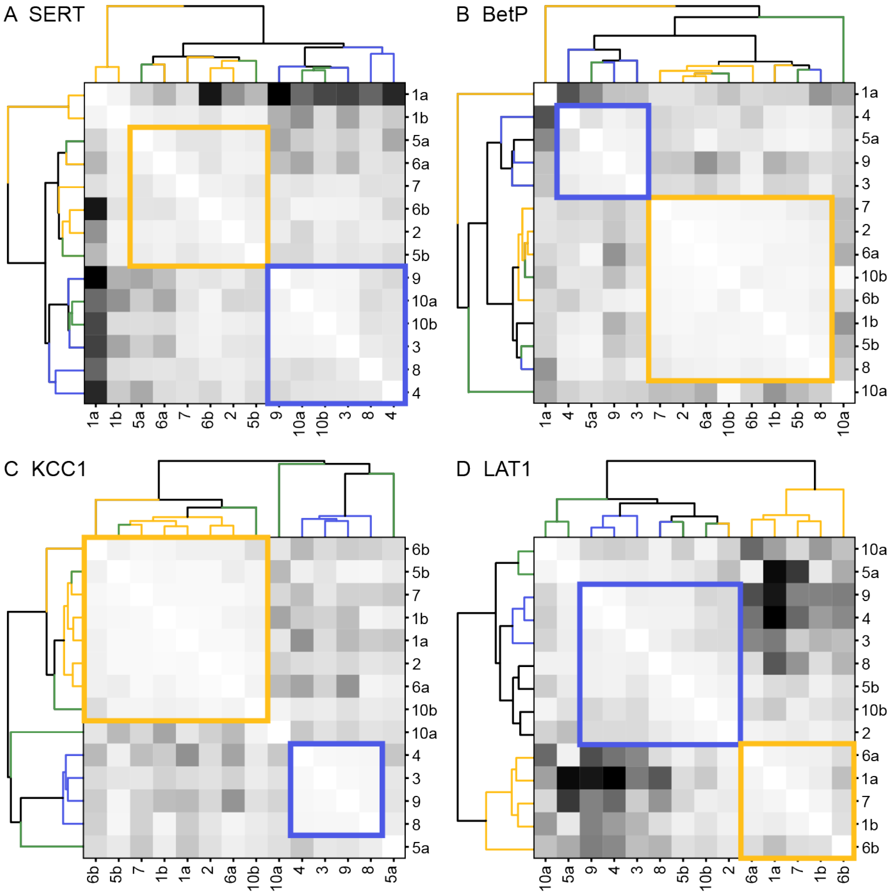


**Figure S6. Clustered hb-DDMs for all available $O_{op}$→$I_{op}$ comparisons show that all seven analyzed LeuT-fold transporters have a rigid hash.**

This figure includes the four clustered hb-DDMs not illustrated Figure 3. The helix labels are colored blue for the hash, yellow for the bundle, and green for the arms. The rigid bodies are highlighted with colored boxes on each matrix, with blue boxes for rigid bodies containing the hash, and yellow boxes for rigid bodies containing the bundle. Only in BetP do the four hash helices (TMs 3, 4, 8, and 9) not all cluster together, with TM8 excluded from the hash cluster and clustering instead with the bundle helices.

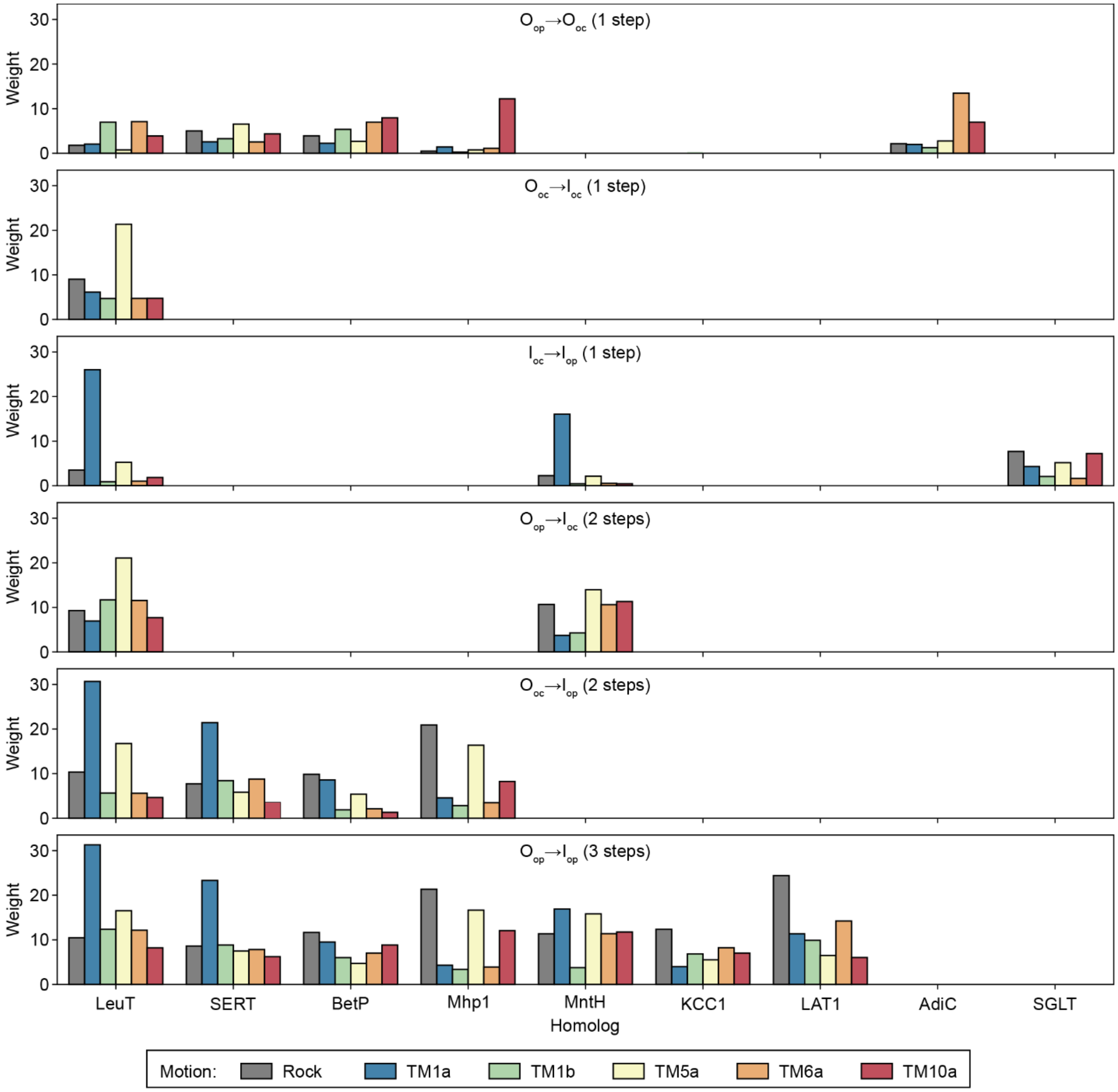


**Figure S7. Sparse PCA weights for all 22 pairwise structural comparisons.**

### Supplementary Materials References


1. Forrest, L.R., Kramer, R., and Ziegler, C. (2011). The structural basis of secondary active transport mechanisms. Biochim Biophys Acta *1807*, 167-188. 10.1016/j.bbabio.2010.10.014.

2. Gao, X., Zhou, L., Jiao, X., Lu, F., Yan, C., Zeng, X., Wang, J., and Shi, Y. (2010). Mechanism of substrate recognition and transport by an amino acid antiporter. Nature *463*, 828-832. 10.1038/nature08741.

3. Coleman, J.A., Yang, D., Zhao, Z., Wen, P.C., Yoshioka, C., Tajkhorshid, E., and Gouaux, E. (2019). Serotonin transporter-ibogaine complexes illuminate mechanisms of inhibition and transport. Nature *569*, 141-145. 10.1038/s41586-019-1135-1.

4. Kazmier, K., Sharma, S., Islam, S.M., Roux, B., and McHaourab, H.S. (2014). Conformational cycle and ion-coupling mechanism of the $Na^+$/hydantoin transporter Mhp1. Proc Natl Acad Sci U S A *111*, 14752-14757. 10.1073/pnas.1410431111.

5. Jeschke, G. (2013). A comparative study of structures and structural transitions of secondary transporters with the LeuT fold. Eur Biophys J *42*, 181-197. 10.1007/s00249-012-0802-z.

6. Kazmier, K., Sharma, S., Quick, M., Islam, S.M., Roux, B., Weinstein, H., Javitch, J.A., and McHaourab, H.S. (2014). Conformational dynamics of ligand-dependent alternating access in LeuT. Nat Struct Mol Biol *21*, 472-479. 10.1038/nsmb.2816.

7. Shimamura, T., Weyand, S., Beckstein, O., Rutherford, N.G., Hadden, J.M., Sharples, D., Sansom, M.S., Iwata, S., Henderson, P.J., and Cameron, A.D. (2010). Molecular basis of alternating access membrane transport by the sodium-hydantoin transporter Mhp1. Science *328*, 470-473. 10.1126/science.1186303.

8. Yan, R., Li, Y., Muller, J., Zhang, Y., Singer, S., Xia, L., Zhong, X., Gertsch, J., Altmann, K.H., and Zhou, Q. (2021). Mechanism of substrate transport and inhibition of the human LAT1-4F2hc amino acid transporter. Cell Discov *7*, 16. 10.1038/s41421-021-00247-4.

9. Bozzi, A.T., Zimanyi, C.M., Nicoludis, J.M., Lee, B.K., Zhang, C.H., and Gaudet, R. (2019). Structures in multiple conformations reveal distinct transition metal and proton pathways in an Nramp transporter. eLife *8*, e41124. 10.7554/eLife.41124.

10. Perez, C., Koshy, C., Yildiz, O., and Ziegler, C. (2012). Alternating-access mechanism in conformationally asymmetric trimers of the betaine transporter BetP. Nature *490*, 126-130. 10.1038/nature11403.

11. Zhao, Y., Shen, J., Wang, Q., Ruiz Munevar, M.J., Vidossich, P., De Vivo, M., Zhou, M., and Cao, E. (2022). Structure of the human cation-chloride cotransport KCC1 in an outward-open state. Proc Natl Acad Sci U S A *119*, e2109083119. 10.1073/pnas.2109083119.

12. Forrest, L.R., and Rudnick, G. (2009). The rocking bundle: a mechanism for ion-coupled solute flux by symmetrical transporters. Physiology (Bethesda) *24*, 377-386. 10.1152/physiol.00030.2009.

13. Bozzi, A.T., Bane, L.B., Weihofen, W.A., McCabe, A.L., Singharoy, A., Chipot, C.J., Schulten, K., and Gaudet, R. (2016). Conserved methionine dictates substrate preference in Nramp-family divalent metal transporters. Proc Natl Acad Sci U S A *113*, 10310-10315. 10.1073/pnas.1607734113.

14. Ehrnstorfer, I.A., Geertsma, E.R., Pardon, E., Steyaert, J., and Dutzler, R. (2014). Crystal structure of a SLC11 (NRAMP) transporter reveals the basis for transition-metal ion transport. Nat Struct Mol Biol *21*, 990-996. 10.1038/nsmb.2904.

15. Koshy, C., Schweikhard, E.S., Gartner, R.M., Perez, C., Yildiz, O., and Ziegler, C. (2013). Structural evidence for functional lipid interactions in the betaine transporter BetP. EMBO J *32*, 3096-3105. 10.1038/emboj.2013.226.

16. Gotfryd, K., Boesen, T., Mortensen, J.S., Khelashvili, G., Quick, M., Terry, D.S., Missel, J.W., LeVine, M.V., Gourdon, P., Blanchard, S.C., et al. (2020). X-ray structure of LeuT in an inward-facing occluded conformation reveals mechanism of substrate release. Nat Commun *11*, 1005. 10.1038/s41467-020-14735-w.

17. Paz, A., Claxton, D.P., Kumar, J.P., Kazmier, K., Bisignano, P., Sharma, S., Nolte, S.A., Liwag, T.M., Nayak, V., Wright, E.M., et al. (2018). Conformational transitions of the sodium-dependent sugar transporter, vSGLT. Proc Natl Acad Sci U S A *115*, E2742-E2751. 10.1073/pnas.1718451115.

18. Simmons, K.J., Jackson, S.M., Brueckner, F., Patching, S.G., Beckstein, O., Ivanova, E., Geng, T., Weyand, S., Drew, D., Lanigan, J., et al. (2014). Molecular mechanism of ligand recognition by membrane transport protein, Mhp1. EMBO J *33*, 1831-1844. 10.15252/embj.201387557.

19. Yamashita, A., Singh, S.K., Kawate, T., Jin, Y., and Gouaux, E. (2005). Crystal structure of a bacterial homologue of $Na^+$/$Cl^-$-dependent neurotransmitter transporters. Nature *437*, 215-223. 10.1038/nature03978.

20. Faham, S., Watanabe, A., Besserer, G.M., Cascio, D., Specht, A., Hirayama, B.A., Wright, E.M., and Abramson, J. (2008). The crystal structure of a sodium galactose transporter reveals mechanistic insights into $Na^+$/sugar symport. Science *321*, 810-814. 10.1126/science.1160406.

21. Bozzi, A.T., Bane, L.B., Zimanyi, C.M., and Gaudet, R. (2019). Unique structural features in an Nramp metal transporter impart substrate-specific proton cotransport and a kinetic bias to favor import. J Gen Physiol *151*, 1413-1429. 10.1085/jgp.201912428.

22. Gupta, K., Donlan, J.A.C., Hopper, J.T.S., Uzdavinys, P., Landreh, M., Struwe, W.B., Drew, D., Baldwin, A.J., Stansfeld, P.J., and Robinson, C.V. (2017). The role of

interfacial lipids in stabilizing membrane protein oligomers. Nature *541*, 421-424. 10.1038/nature20820.

23. Kilic, F., and Rudnick, G. (2000). Oligomerization of serotonin transporter and its functional consequences. Proc Natl Acad Sci U S A *97*, 3106-3111. 10.1073/pnas.97.7.3106.

24. Perez, C., Khafizov, K., Forrest, L.R., Kramer, R., and Ziegler, C. (2011). The role of trimerization in the osmoregulated betaine transporter BetP. EMBO Rep *12*, 804-810. 10.1038/embor.2011.102.

25. Weyand, S., Shimamura, T., Yajima, S., Suzuki, S., Mirza, O., Krusong, K., Carpenter, E.P., Rutherford, N.G., Hadden, J.M., O'Reilly, J., et al. (2008). Structure and molecular mechanism of a nucleobase-cation-symport-1 family transporter. Science *322*, 709-713. 10.1126/science.1164440.

26. Verrey, F., Closs, E.I., Wagner, C.A., Palacin, M., Endou, H., and Kanai, Y. (2004). CATs and HATs: the SLC7 family of amino acid transporters. Pflugers Arch *447*, 532-542. 10.1007/s00424-003-1086-z.

27. Casula, S., Zolotarev, A.S., Stuart-Tilley, A.K., Wilhelm, S., Shmukler, B.E., Brugnara, C., and Alper, S.L. (2009). Chemical crosslinking studies with the mouse Kcc1 K-Cl cotransporter. Blood Cells Mol Dis *42*, 233-240. 10.1016/j.bcmd.2009.01.021.
